# HIV-1 Integration Site Determines the Transcriptional Fate and Persistence of Integrated Proviruses

**DOI:** 10.64898/2025.12.26.696579

**Authors:** Virender Kumar Pal, Ali Danesh, Marie Canis, Thomas R. Dilling, Itzayana G. Miller, Tan Thinh Huynh, Dennis C. Copertino, Douglas Barrows, Thomas Carroll, Theodora Hatziioannou, R. Brad Jones, Guinevere Q. Lee, Frauke Muecksch, Paul D. Bieniasz

**Affiliations:** Laboratory of Retrovirology, The Rockefeller University, New York; Division of Infectious Diseases, Department of Medicine, Weill Cornell Medical College, New York; Bioinformatics Resource Center, The Rockefeller University, New York; Department of Infectious Diseases, Virology, Heidelberg University, Heidelberg, Germany; Howard Hughes Medical Institute, The Rockefeller University, New York

## Abstract

The mechanisms by which latent HIV-1 reservoirs persist during antiretroviral therapy is incompletely understood. Here, we derive a model system to measure clonal expansion and viral latency in which populations of human memory CD4^+^ T cells, each bearing a single transcriptionally active HIV-1 provirus are engrafted into immunodeficient mice. Over ∼2 months *in vivo*, clonal expansion and the establishment of latency occurred in subsets of engrafted infected cells. Clonal expansion *in vivo* was driven by T-cell receptor identity, but not by proviral insertional mutagenesis. The integration sites of proviruses that became latent *in vivo* were enriched on chromosome 19, in intergenic and centromeric satellite regions, and genes whose expression is atypically low. Pre-existing repressive epigenetic features were associated with latency for subsets of proviruses. Our findings suggest a confluency of genomic and epigenomic factors predispose certain genomic locations, including ZNF genes, to host proviruses that constitute the latent reservoir.

## Introduction

During antiretroviral therapy (ART) of HIV-1 infection, a reservoir of CD4^+^ T-cells harboring intact HIV-1 proviruses persists and comprises a major barrier to achieving cures. These reservoir-harboring cells undergo clonal expansion, evade host immune responses, and proviruses therein provide a source of viral rebound upon ART interruption^1–6^. Understanding the mechanisms underlying the establishment and maintenance of the HIV-1 proviral reservoir may provide an opportunity to purge the reservoir and remove the need for lifelong ART. However, there are a number of obstacles to achieving a detailed understanding of proviral persistence. The extreme rarity of cells harboring intact proviruses *in vivo,* typically ∼one in a million CD4^+^ T cells, poses a significant challenge to the investigation of the mechanisms that enable their persistence^7,8^. Moreover, the intact proviral DNA that is intrinsically capable of initiating viral rebound, is partly obscured by the existence of a larger amount of integrated viral DNA that is not intact, but becomes comparatively abundant due to the selective survival of T-cell clones that harbor defective, non-expressed proviruses ^9–12^.

Several factors may govern the establishment and maintenance of the HIV-1 reservoir. A key mechanism of persistence is proviral latency, which protects infected cells from viral cytopathicity and immune clearance. Whether a provirus adopts a state of latency versus active transcription can be impacted by the physiological state of the infected cell, which influences the levels of trans-acting factors such as NF-κB and NFAT, that are activated during T-cell activation and link cellular activation to HIV-1 transcription^13,14^. Additionally, immune evasion mechanisms, such as impaired antigen presentation or resistance to cytotoxic T lymphocyte (CTL) killing, may further promote the survival and expansion of reservoir cells during periods of activation ^15–19^.

The site of proviral integration into host DNA may influence reservoir persistence in two key and distinct ways. First, the genomic context of the integration site may affect the accessibility of proviruses to the transcriptional machinery and thus impact the propensity of a provirus to establish and maintain latency. Second, by acting as an insertional mutagen, HIV-1 integration in or near genes controlling cell proliferation may drive clonal expansion ^4,20–29^. HIV-1 favors integration within actively transcribed genes, via the interactions of viral capsid and integrase proteins with CPSF-6 and LEDGF/p75, host proteins that are associated with areas of the genome, or individual genes, that are actively transcribing ^30–35^. In HIV-1-infected expanded cell clones from ART-treated individuals, viral integration sites are often found in a small number of genes whose function is related to cell survival and proliferation, suggesting that clonal expansion might be driven in part by viral integration. However, these clones typically harbor defective proviruses, and the mechanism by which these insertions drive clonal expansion, and whether they contribute to intact proviral reservoir persistence, remains unclear ^5,27,36^. In contrast, recent studies have shown that expanded clones of cells from infected individuals on long-term ART that carry intact proviruses, do so predominantly with proviral integration sites in regions comprising gene deserts, intergenic, centromeric satellite DNA, and also, inexplicably, within zinc finger (ZNF) genes ^23,29,37^.

Studies of integrated HIV-1 proviruses in cell lines have suggested the importance of the genomic context in latency and its reversal ^38,39^. However, currently available *in vitro* models for studying HIV-1 latency in primary CD4^+^ T cells ^40–44^ do not allow the evaluation of the relative contributions of integration site versus the intrinsic physiological properties of T cell clones harboring proviruses. Long-term *ex vivo* cell culture models in which primary CD4^+^ T cells from ART-treated individuals have been used to model latency in an authentic cell context, but cannot evaluate cell clone-specific effects ^45,46^. Thus, there is an unmet need for laboratory methods capable of identifying individual HIV-1-infected cell clones, tracking their long-term clonal expansion, and assessing the effects of proviral integration site on latency in a clone-specific manner.

In this study, we present a novel ‘HIV-1 latency mouse’ (HIVLM) model that overcomes some of the limitations of current *in vitro* and *ex vivo* models of HIV-1 latency. We use the model to determine the impact of proviral integration sites on clonal expansion and latency in HIV-1 infected human memory CD4^+^ T cells. Combining the HIVLM model with a method for efficient detection of integration sites and quantification of clonal expansion (PRISM-seq) ^47^, we determined how host-specific genomic and epigenomic features associated with each distinct HIV-1 integration site affect clonal survival and proviral expression in the absence of immunological or anti-viral drug selection pressures. We find that clonal expansion of infected cells in the HIVLM model was linked to the T cell receptor (TCR) sequence of engrafted cells. We find no evidence that clonal expansion is driven by insertional mutagenesis associated with provirus integration into specific genes. In contrast, the proviral genomic location and epigenetic status proximal to the integration site are key factors distinguishing latent from active proviruses. Latent proviruses were found at elevated frequency in intergenic regions and associated with heterochromatin. In line with previous clinical studies, latent proviruses in the HIVLM model were also found at elevated frequency in ZNF genes. Notably, our analysis reveals that the increased number of latent proviruses in ZNF genes was associated with their abundance, location, low expression levels, and heightened repressive epigenetic state.

## Results

### Long-term *in vivo* survival of HIV-1 infected human memory CD4^+^ T cells in the HIVLM model

To track HIV-1 latency, we developed a single-cycle HIV-1 reporter virus construct, termed V1/SBP-GFP, that expresses both GFP, and a cell surface streptavidin binding peptide (SBP) fused to low-affinity nerve growth factor receptor. These reporters, expressed in place of Nef under the control of the authentic HIV-1 promoter and splicing signals, could be used to both monitor HIV-1 gene expression and as an affinity tag for one-step positive or negative selection of cells with active or latent proviruses, using streptavidin-conjugated magnetic beads.

The suitability of primary human CD4^+^ T cells for the study of HIV-1 latency over long timespans is limited by their relatively short lifespan in cell culture. Following activation and infection of freshly isolated CD4^+^ T cells, these cells can be maintained in culture for only a few weeks, with a need for re-stimulation to propagate them further. To enable the investigation of HIV-1 latency and persistence in primary human CD4^+^ T cells over longer time periods, we adapted a previously described participant-derived xenograft (PDX) model of HIV-1 infection^48^. In this model, NSG mice are engrafted with the memory (CD45RO^+^) subset of CD4^+^ T cells, which avoids the severe graft versus host disease that rapidly occurs in mice engrafted with total human CD4^+^ T cells^48^. For mouse engraftment, memory CD4^+^ T cells were enriched from leukapheresis-derived PBMCs of HIV-1-negative donors via negative selection (Fig. S1a). Cells were activated for 48 hours by CD3/CD28-stimulation, resulting in >80% in CD69^+^ cells (Fig. S1b). Activated cells were infected, and approximately 4×10^9^ cells were subjected to streptavidin-bead-based magnetic cell sorting to isolate V1/SBP-GFP-expressing memory CD4^+^ T cells. Infection was done at low multiplicity so that after positive selection, each cell is presumed to carry a single transcriptionally active provirus at a unique integration site (Fig.1a). The sorting procedure yielded a >90 % pure cell population of V1/SBP-GFP-expressing cells (Fig. S1c). V1/SBP-GFP expression was confirmed two days after sorting, and cells were cryopreserved in aliquots for later engraftment into mice (Fig. S1d).

**Figure 1.**
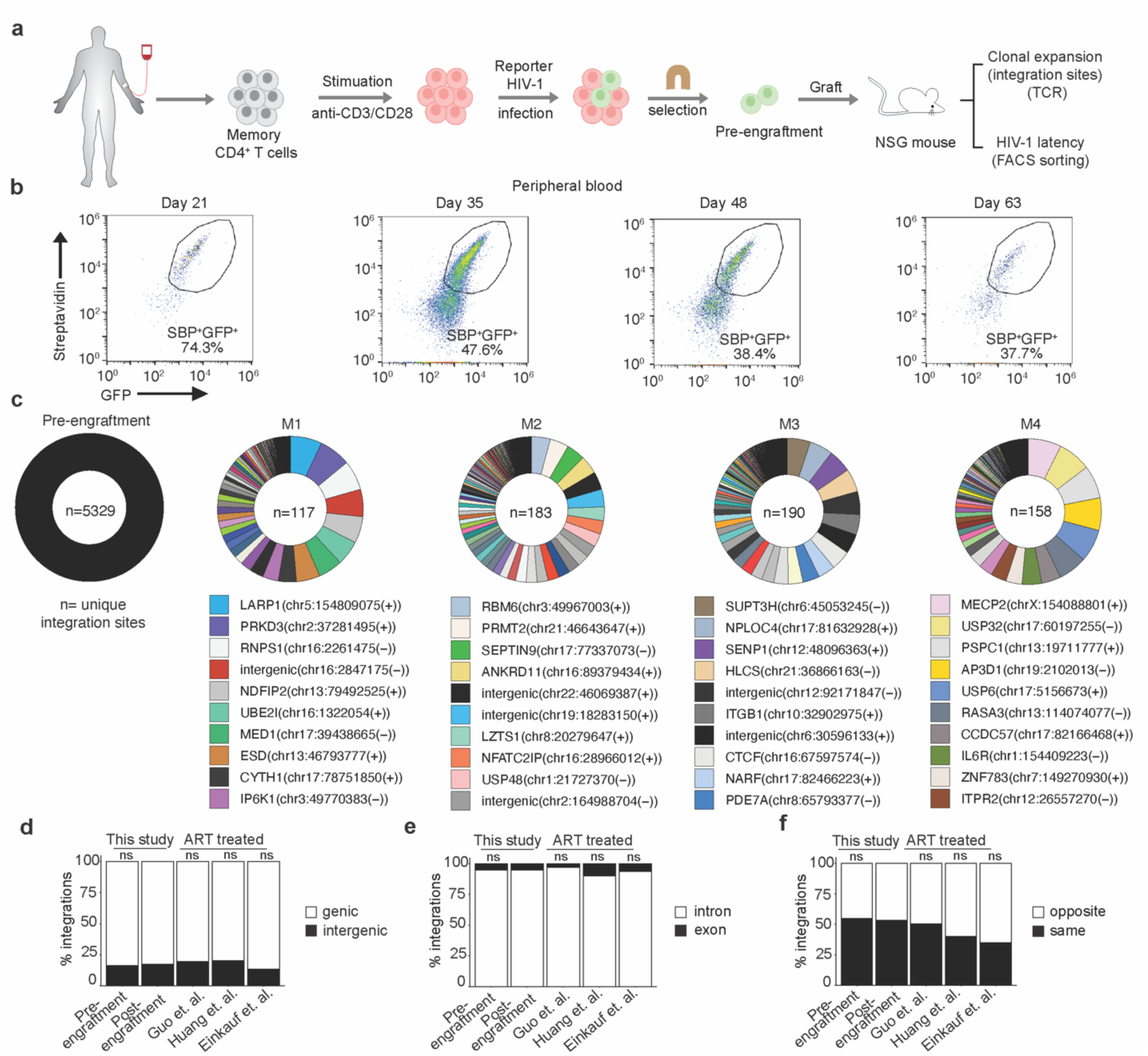
Tracking clonal expansion of HIV-1 infected human memory CD4^+^ T cells *in vivo* in HIVLM model. **a)** Isolation of human CD4^+^ memory T cells, reporter HIV-1 (V1/SPB-GFP) infection, selection of infected cells, and engraftment in NSG mice. **(b)** Flow cytometry-based temporal analysis of GFP/SBP expression in engrafted human cells in the peripheral blood of a typical mouse. **(c)** Circos plots showing the relative expansion of HIV-1 infected cell clones in the pre- and post-engraftment (spleen) cell populations. Each clone is color-coded according to its proviral integration site, and the area of the pie represents the relative extent of clonal expansion, as measured by the number of wells in which that integration site was detected. Each integration site is labelled as the name of the gene with integrated provirus or as ‘intergenic’ if integration was outside of the gene, followed by the chromosome name, the genomic coordinate, and the host DNA strand (positive or negative). The n value represents the total number of unique integration sites identified in an individual pre- or post-engraftment (spleen) cell population. **(d)** Comparison of genic and intergenic distribution of unique integration sites in pre- or post-engraftment (spleen) cell populations, and a previously published clinical datasets^11,23,50^. **(e, f)** Comparison of the proportion of genic integrations present in introns versus exons (**e**), and in same or opposite orientation as host transcription (**f**). The p-values for differences compared to post-engraftment (spleen) cell populations were determined by Fisher’s exact test; ns = non-significant.

The transcriptionally active HIV-1 provirus-containing (SBP^+^GFP^+^) human memory CD4^+^ T cells were engrafted into NSG mice by tail vein injection (∼5-7×10⁶ cells per mouse), and successful engraftment was confirmed by the subsequent appearance of human CD45⁺ cells in peripheral blood (Fig. 1b and S1e). Engrafted human CD4^+^ T cells were detected in all mice, with ∼10-100 human CD4^+^ T cells per mm^³^ blood at 35 days after engraftment (Fig. S1f). At this time point, ∼40 to 65% of the human CD45⁺ cells were SBP⁺GFP⁺, and a time-dependent loss of SBP/GFP reporter expression ensued, (Fig. 1b and S1g). The loss of SBP/GFP expression in human CD4^+^ T cells was observed in multiple tissues, including spleen, lung, and bone marrow, months after engraftment (Fig. S1h-j). To determine whether the loss of SBP/GFP expression over time reflected the establishment of latency or the selective outgrowth of contaminating uninfected cells, we determined the *psi-gag*, *env*, and human RPP30 DNA copy numbers in genomic DNA extracted from mouse blood and tissues by digital droplet PCR (ddPCR) using an intact proviral DNA assay (IPDA)^49^. Detection of approximately one *psi-gag* and *env* copy per human cell demonstrates the long-term survival of HIV-1 infected memory CD4^+^ T cells and shows that most human cells in mouse spleens contained a reporter provirus (Fig. S1k), despite the low number of GFP-expressing cells. The observed loss of V1/SBP-GFP expression is therefore not caused by outgrowth of contaminating uninfected cells, but primarily by the transition to latency by a subset of proviruses in infected cells, during the months after engraftment in the HIVLM model.

### Clonal expansion of HIV-1 infected human cells and proviral integration site features in the HIVLM model

To determine the distribution of integration sites in engrafted HIV-1 infected human cells, we employed a method, termed PRISM-seq, in which a DNA sample is divided into 96 replicates, each containing an estimated 50 proviruses, as measured by ddPCR, prior to whole genome amplification and integration site determination. PRISM-seq enables the identification of >80% of the integration sites in a sample, and the fraction of the 96 replicates in which a given integration site is found is a measure of the clonal expansion of the T cell containing an HIV-1 provirus with that integration site ^47^. Using this approach, we simultaneously determined HIV-1 integration site identity and the extent of clonal expansion for each HIV-1-infected memory CD4^+^ T cell harboring a given integration site, for the pre-engraftment cell population, and for cells recovered from 8 mouse spleens >2 months after engraftment of cells from the same donor. Post-engraftment spleen samples were either left unsorted or were sorted into active and latent cell populations, based on SBP/GFP expression, before HIV-1 integration site determination (Fig. S2a,b,c and S3a,b). We identified 5239 and 3152 unique integration sites in pre-engraftment and post-engraftment cell populations, respectively (Fig. 1c, S2a,b and S3a). For 57% to 84% of proviruses, we were able to capture both the 5’ and 3’ LTR-host junction during integration sites determination (Fig. S2b and S3a), indicating that our overall integration site detection was efficient, and suggesting (along with the SBP/GFP and IPDA measurements conducted on the pre- and post-engraftment samples) that the majority of proviruses were intact.

Clonal expansion of the HIV-1-infected cells in the pre-engraftment sample was minimal, with 3.1% of the 5239 integration sites detected in more than one replicate. This result was approximately as expected given that the cells were sampled 4 days after *in vitro* infection (Fig. S2a). Conversely, at > 2 months after mouse engraftment, clonal expansion of HIV-1-infected cells was extensive, with 8% to 60% of integration site detected multiple times in 96 replicates, and with a few HIV-1 integration sites appearing in all 96 replicates of the PRISM-seq assay (representing a frequency of > 1 in 50 cells and thus exceeding the dynamic range of our clonal expansion measurement) (Fig. 1c, S2a and S3a). Consistent with previously published data, HIV-1 integrations were biased toward genic rather than intergenic regions in the pre-engraftment sample ^50^. Moreover, there was no clear difference between pre- and post-engraftment samples in genic versus intergenic distribution of proviral integration sites (Fig. 1d, S2c, and S3b), The proportion of proviruses integrated in genic or intergenic regions in cells recovered from mice was similar to that reported for ART-treated individuals^11,23,51^ (Fig. 1d). Most genic integration sites were in introns, as expected (Fig. 1e) and approximately equivalent numbers of proviruses were integrated in the same versus the opposite orientation with respect to transcription of the gene in which the integration site was found (Fig. 1f). Overall, these results suggest that the genomic features associated with HIV-1 integration sites obtained pre-engraftment, after *in vitro* infection of primary CD4^+^ T-cells, and also after 2 months of engraftment in the HIVLM model were consistent with patterns observed clinically in ART-treated individuals.

### Frequent integration of HIV-1 into specific host genes does not predict clonal expansion

Because HIV-1 has the potential to act as an insertional mutagen that might drive cell proliferation in humans or the mouse grafts, we next examined whether HIV-1 integration into specific host genes predicted the extent of expansion of the respective infected human cell clone during the ∼ 2 month engraftment period. We identified numerous genes, such as NPLOC4, PACS1, NFATC3, NOSIP, NEAT1, ASH1L, and KDM2A, in which multiple distinct HIV-1 integrations were found in multiple mice, following 2 months of engraftment (Fig. 2a). However, the appearance of multiple integrated proviruses in a given gene across several engrafted cell populations did not predict the degree of expansion of the respective cell clones containing that integration site. A possible exception to this conclusion was NPLOC4, in which one out of 10 cell clones was highly expanded in one mouse (Fig. 2a). A better predictor of the frequency with which proviral integrations were found in a given gene post engraftment, was the number of integration sites found in that same gene in the pre-engraftment population (Fig. 2b). The genes with increased frequency of integration were clustered near previously identified *in vitro* infection defined integration ‘hot spots’^52^, for example on chromosome 11, 16, and 17 in both pre- and post-engraftment cell populations (Fig. S4). As reported previously for *in vitro* HIV-1 infections, increased gene density correlated with increased integration frequency in both pre- and post-engraftment cell populations^53–55^ (Fig. S4), The genes with frequent integration in our pre- and post-engraftment datasets also overlap with the previously described “recurrent integration genes” (RIGs)^52,56,57^ (Fig. S5a and S5b).

**Figure 2.**
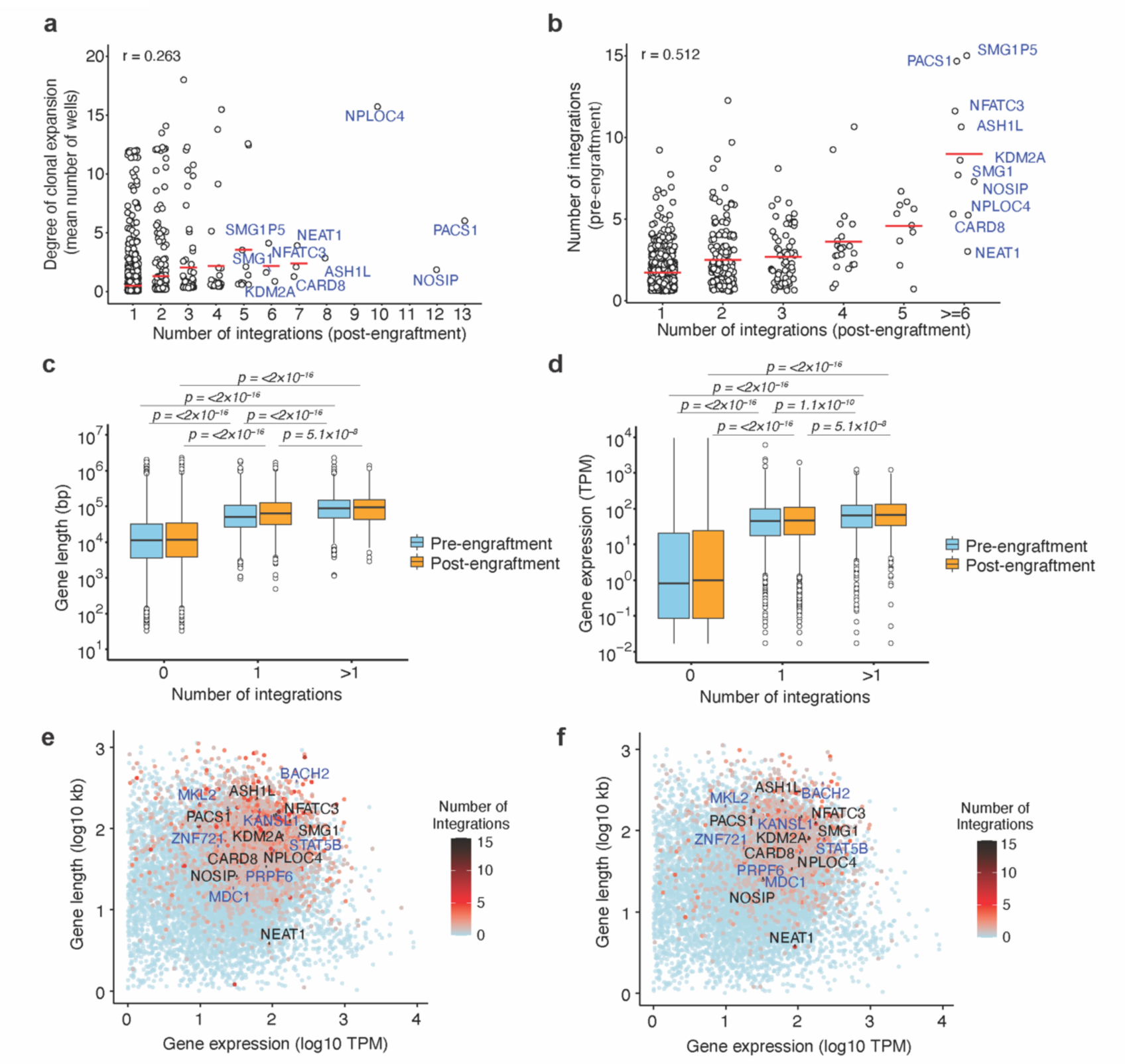
Identification of genes with frequent HIV-1 integration and their association with clonal expansion. **(a)** Extent of clonal expansion, as measured by the mean number of wells in which a unique integration site in a given gene was detected, plotted against the number of unique integration sites in a given gene detected post-engraftment. Each data point represents a gene with at least one integrant. **(b)** The number of unique integrations in a given gene in the pre-engraftment cell population plotted against the number of unique integration sites detected in that same gene 2 months post-engraftment. For **a** and **b**, data from four mice whose cells were not sorted is pooled. **(c, d)** Box plot of gene lengths (**c**) and level of expression (**d**) for genes with 0, 1, or >1 integrations. The plot shows median, interquartile ranges, and minimum/maximum values. P values were determined by pairwise Wilcoxon rank-sum tests with Bonferroni correction for multiple comparisons. **(e, f)** Gene length and expression level for genes without or with a varying number of integrated proviruses as indicated by the color scale in (e) pre-and (f) post-engraftment samples.

Comparison of gene length and gene expression levels, determined based on previously published RNA-seq data from activated central memory CD4^+^ T cells^58^, revealed that both characteristics significantly correlated with an increased number of HIV-1 integrations in both pre- and post-engraftment samples (Fig. 2c and 2d, respectively). Genes such as NPLOC4, PACS1, NFATC3, NOSIP, NEAT1, ASH1L, and KDM2A may therefore be frequent targets of integration because they are large genes and/or are highly expressed in memory CD4^+^ T cells (Fig. 2e,f). CPSF6 and LEDGF bias HIV-1 integration toward gene bodies and to nuclear speckles^32,34,54^. Using published TSA-seq and DamID datasets for speckle-associated domains (SPAD) and lamina-associated domains (LAD)^59^, we also found enrichment of proviral integration sites in SPAD (∼75%) as compared to LAD (Fig. S5c) in both pre- and post-engraftment populations. Indeed, genes with >1 proviral integration were more likely to be found in SPADs compared to genes without proviruses (Fig. S5d and S5e). In ART-treated individuals, proviral integration sites in expanded clones have been reported near host genes associated with cell growth and proliferation, suggesting a possible effect of insertional mutagenesis on proliferation and survival of provirus-containing cell clones^4,5^. However, comparative ontology analysis of genes with HIV-1 integrations in pre- and post-engraftment cell populations showed a similar distribution of integration sites in genes associated with various functions, including epigenetic regulation of gene expression, lymphocyte differentiation, and mitotic cell cycle regulation (Fig. S5f). Additionally, in pre- and post-engraftment cells, we observed no enrichment or depletion of integration sites in essential versus non-essential genes in expanded clones as compared to non-expanded clones, irrespective of whether they were in RIGs or not^60^ (Fig. S5g). Overall, the distribution of HIV-1 integration sites among human genes was similar for *in vitro* infected memory CD4^+^ T-cells (pre-engraftment) and among clonally expanded HIV-1 infected CD4^+^ T cells in mice (post-engraftment). Because the pattern of integration sites did not differ in these two populations, these results suggest an absence of selection pressure for proliferation and survival that is driven by HIV-1 integration into specific genes. Rather, specific genes were found to harbor proviruses in engrafted cells, including those that were clonally expanded, because they are more frequent targets for provirus integration, due to their size, expression level, and nuclear location, not because of putative effects of proviral integration on CD4^+^ T-cell survival or proliferation.

### T cell receptor (TCR) association with clonal expansion

Mature T cells harbor uniquely rearranged TCR genes, whose sequence can be used to track the expansion of individual T cell clones. While it was not possible to link individual TCR clonotypes to a given HIV-1 integration site in our analyses, we determined complementarity-determining region 3 (CDR3) TCR sequences in human cells pre- and post-engraftment, to assess clonal expansion (Fig. S6a,b). We found hundreds of unique TCR in each engrafted mouse (mean = 300, range 111 to 638), similar to the predicted number of independent T-cell clones recovered after engraftment inferred by HIV-1 integration site mapping (Fig. S6b and Fig. 1c). The number of unique T cell clones recovered from individual mice at 2 months post-engraftment (mean = 300) was lower than the number of T cell clones recovered from the pre-engraftment sample (n = 38,621), indicating a bottleneck for cell survival or selective clonal expansion following engraftment (Fig. S6b). While each mouse harbored a distinct TCR repertoire, and most TCR sequences were recovered from only one mouse, certain TCR variants were detected in 2 or more mice post-engraftment for both HIV-1 infected and uninfected engraftments (Fig. 3a and 3b). Indeed, certain T cell clones were expanded similarly across multiple mice, whether or not the cells were HIV-1 infected prior to engraftment (Fig. 3c-g). For example, a T-cell clone harboring the CDR3 sequence CASSYGGAGNYGYTF was similarly expanded in HIV-1 infected and uninfected grafts (Fig. 3c-f). In that case, the clone-CASSYGGAGNYGYTF was associated with an abundant clone in the pre-engraftment sample (Fig. 3g, S6c). However, apparently selective growth of some T cell clones that were either less abundant or not detected in the pre-engraftment sample was evident in one or more engrafted mice (Fig. 3g and S6c). Indeed, while there was no correlation between TCR sequence abundance in the pre-engraftment population and it abundance post-engraftment, the mean number of times a given TCR sequence was recovered from individual mice was correlated with the number of mice from which that clonotype was recovered (Fig 3c,d). These findings indicate that the extent of expansion of HIV-infected and uninfected cell clones post-engraftment is independent of their abundance pre-engraftment and predicted to some extent by their TCR sequence. Xenoreactivity thus likely plays a role in clone selection, a notion that is supported by the frequent occurrence of graft vs host disease in human T-cell xenograft models^61,62^, where the memory CD4^+^T-cells are not purified before engraftment. While our data do not identify specific drivers for the differential expansion of human memory CD4^+^ T cell clones in the HIVLM model, we can nevertheless conclude that a diverse set of memory CD4^+^ T-cells are expanded post-engraftment, predicted in part by TCR sequence and not by HIV-1 integration site. Thus, for the majority of integration sites, HIV-1 proviruses act as passengers, and not as mutagenic drivers during clonal expansion of infected cells in our model, and likely in the majority of cases in humans.

**Figure 3.**
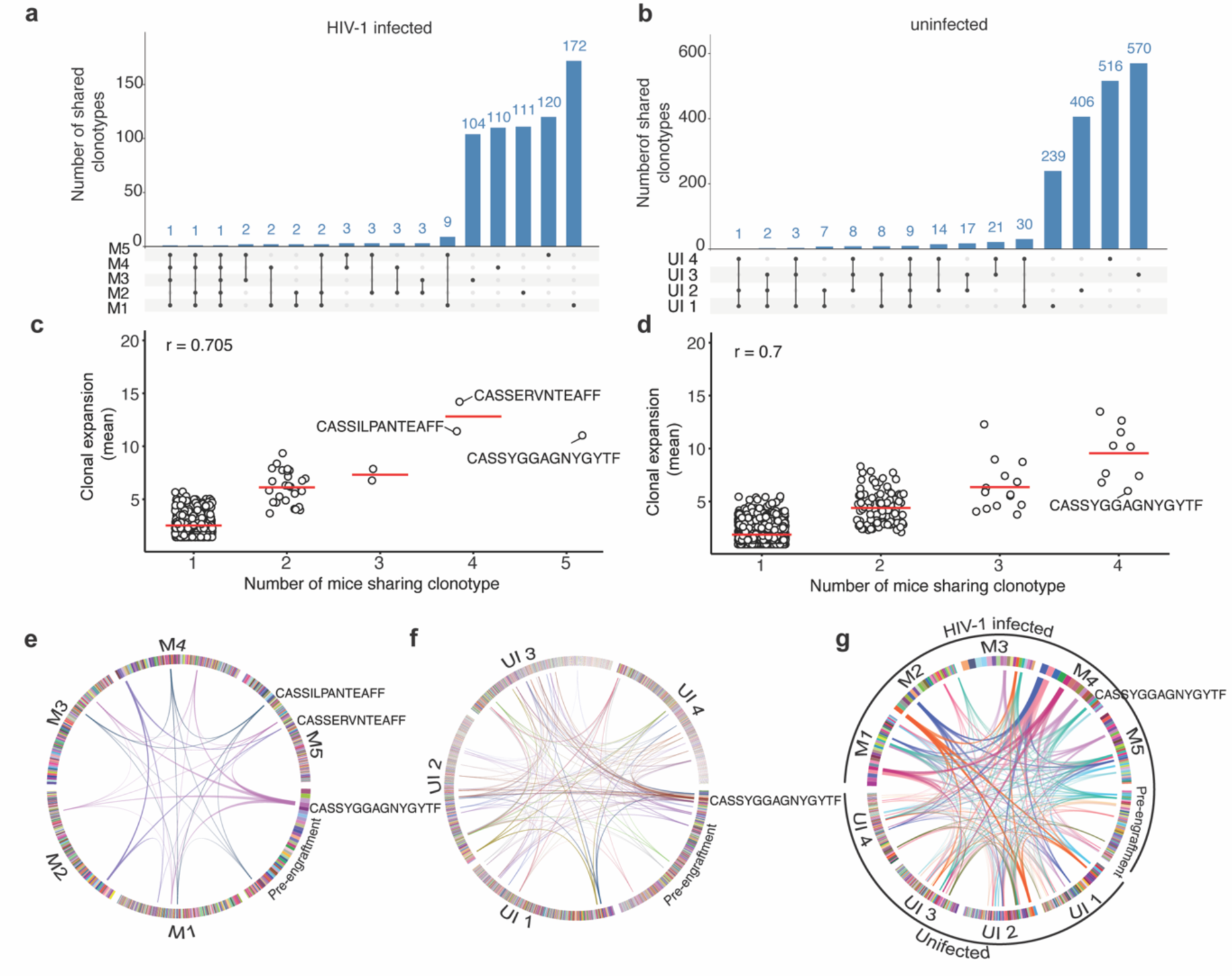
TCR analysis of HIV-1 infected and uninfected human memory CD4^+^ T cells post-engraftment in mice. **(a and b)** Upset plots comparing unique T cell clones retrieved from multiple mice engrafted with HIV-1 infected (a) or uninfected (b) human cells from the same donor. **(c** and **d)** Correlation between the extent of clonal expansion and the number of mice from which individual T cell clonotypes were retrieved after engraftment of HIV-1 infected (c) or uninfected (d) human cells. The r value denotes the Pearson correlation coefficient value. **(e** and **f)** Circos plots showing the extent of clonal expansion of each T cell clone in pre-engraftment and post-engraftment in individual mouse samples. Arcs mark clones shared by a minimum of 4 mice for HIV-1 infected (e) and uninfected (f) grafts. Each T cell clone is color-coded and shown as boxes at the circumference of the circos plot for the respective sample. The T cell clones were arranged clockwise from most expanded to least expanded. **(g)** Circos plot showing T cell clones identified in pre-engraftment, and uninfected, HIV-1-infected post engraftment populations for a shared by a minimum of 4 mice.

### Genomic locations of transcriptionally active and latent proviruses

The genomic context in which an HIV-1 provirus resides in an infected cell *in vivo* could affect its transcription status and its propensity to transition to latency^11,28^. Infected human cells harvested from the HIVLM model >2 months post-engraftment were sorted into populations with transcriptionally active (SBP^+^GFP^+^) or latent (SBP^-^GFP^-^) proviruses. The low-template, single-molecule detection efficiency of PRISM-seq, combined with multiple displacement amplification performed directly on cells isolated from sorted populations, enabled reliable and comprehensive determination of HIV-1 integration sites in both transcriptionally active and latent provirus-harboring cell populations (Fig. S7a and Fig. S7b). This analysis identified 1,347 integration sites associated with transcriptionally active proviruses, 863 sites associated with latent proviruses, and 140 sites associated with both transcriptionally active and latent proviruses (Fig. 4a and Fig. S7c). Note that 2 and 12 integration sites with active and latent proviruses, respectively, were found in more than one mouse and are merged for downstream analysis. Clonal expansion, that is detection in multiple replicate wells was evident for 139 integration sites with only transcriptionally active proviruses (mean clonal size = 7.2 wells, range = 2-93) and 90 integration sites with only latent proviruses (mean clonal size = 2.4 wells, range = 2-10) (Fig 4a). In the comparative analyses that follow, we considered either (i) all detected integration sites or (ii) only the integration sites associated with expanded clones. Restricting the analysis to expanded clones reduced the number of sites that were considered but allowed more confident assignment of active or latent status, as each site was observed multiple times. Additionally, to increase the robustness of our comparisons, we combined all the active and latent integration sites retrieved from four individual mice (Fig. 4a).

**Figure 4.**
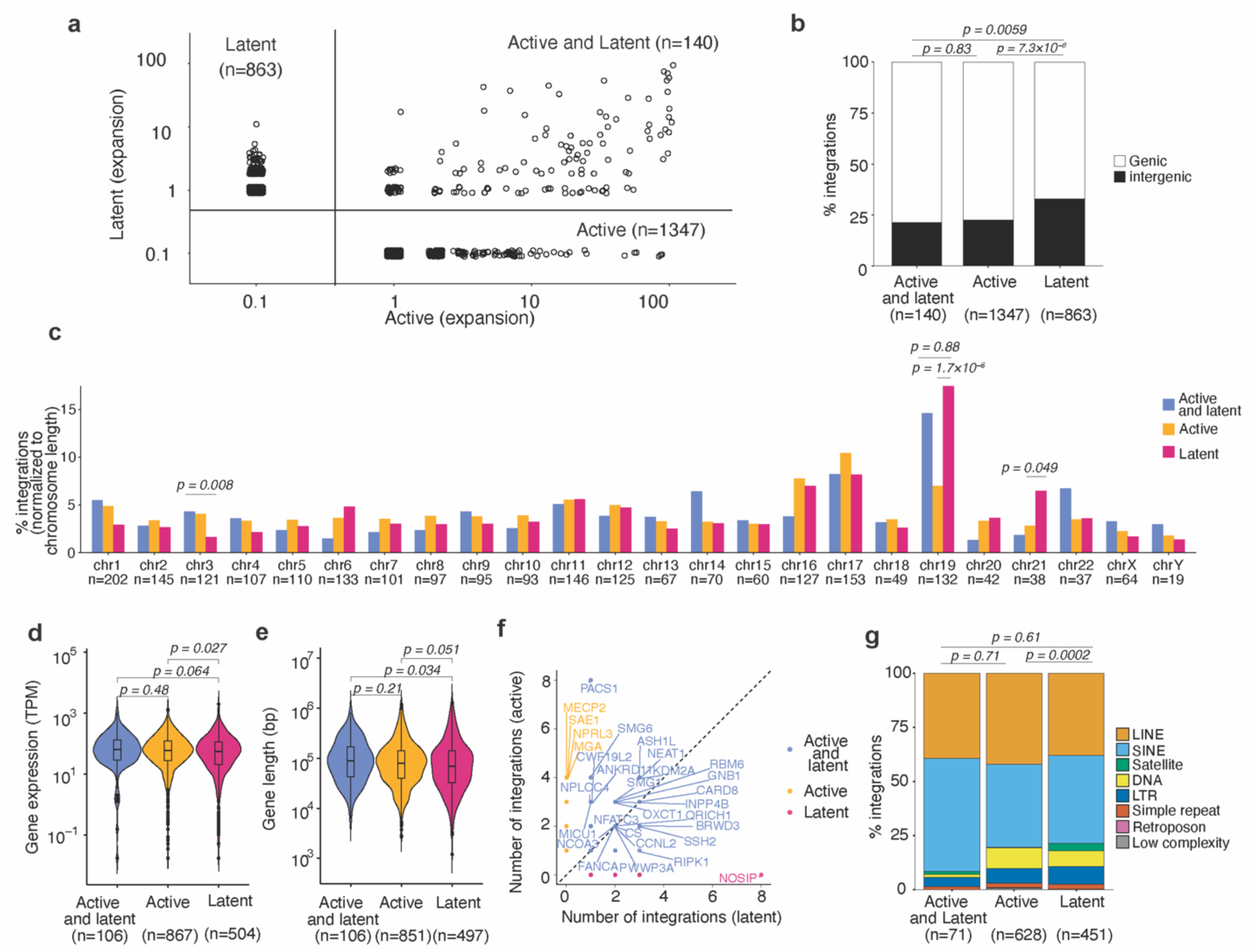
Genomic features associated with active and latent proviral integration sites. **(a)** Expansion of HIV-1-infected cell clones harboring only active, only latent, or a mixture of cells with active or latent proviruses. Axis values represent the number of wells in which an integrant was detected, integrant that were not detected are assigned a value of 0.1 for representation on a logarithmic scale. **(b)** Percentage of genic and intergenic integration site obtained in active, latent, or mixed active and latent cell clones. **(c)** Distribution of active, latent, or active and latent proviruses integration sites on human chromosomes. **(d)** Expression level of genes with only active proviruses, only latent proviruses, or both active and latent proviruses. **(e)** Length of genes with only active proviruses, only latent proviruses, or both active and latent proviruses. **(f)** Number of distinct integration sites in genes with only active proviruses, only latent proviruses, or both active and latent proviruses. The color of the dots and labels indicates if the proviruses in a given gene was found to be active, latent, or exhibited both phenotypes. **(g)** Percentage of integration sites in distinct chromosomal repeat regions in the human genome for the active, latent, or active and latent proviruses. P-values were calculated using Fisher’s exact test (**b** and **c**), Wilcoxon rank-sum test with Bonferroni correction for multiple comparisons (**d** and **e**), and two proportions z-test to compare the proviruses present satellite repeat-rich region between the active and latent populations (**g**).

Latent proviruses were found to be more frequently integrated in intergenic regions as compared to active proviruses, or proviruses exhibiting both active and latent states (Fig. 4b). This differential distribution of integration sites was especially evident when expanded clones were analyzed separately from non-expanded clones (Fig. S7d and S7e). Notably, integration sites harboring latent proviruses, or a mixture of active and latent proviruses, were more likely to be found on chromosome 19 (*p* = 1.7×10^-6^) and 21 (*p* = 0.049) as compared to integration sites of active proviruses (Fig. 4c). This conclusion held for all integration sites found in expanded and non-expanded clones, but was particularly evident for integration sites found in the expanded clones (Fig. S8a-b, S9 and S10).

To ascertain whether the expression levels of a gene was associated with the propensity of proviruses integrated therein to establish latency, we analyzed integration site and host gene expression data in two ways: First, we assigned each gene to a category, determined by whether proviruses integrated therein were active, latent or a mixture of both active and latent, regardless of the number of integration sites in that gene. Second, we assigned each individual genic integration site to an active, latent, or both active and latent category, regardless of the viral phenotype associated with other integration sites in that gene. In the first analysis, human genes with only latent proviruses had lower expression levels than those with only active proviruses (*p = 0.027*) (Fig. 4d and S11a-b). In the second analysis, integration sites with latent proviruses were found in genes with lower expression than integration sites with active proviruses (*p = 0.0094*) (Fig. S11c). The magnitude of these differences was modest but was increased when integration sites in expanded clones were considered separately (Fig. S11d-e). Thus, integration into human genes that have lower expression levels increases the probability of generating a latent provirus. We did not find differences in gene length for human genes with only latent proviruses as compared to active proviruses (Fig. 4e). Notably, most genes in which multiple independent integrations occurred had integration sites with mixtures of both active and latent proviruses. However, some genes exhibited a striking skew in proviral transcriptional activity for the multiple proviruses integrated therein. For example, 7/8 integration sites in PACS1 had active proviruses and one provirus was found in both active and latent populations (Fig. 4f). In contrast, all 8 integration sites in the NOSIP gene harbored only latent proviruses (Fig. 4f, S11f-g). For comparison, the overall fraction of genes with integration sites harboring only active proviruses was 64.3% (number of genes with only active proviral integration sites divided by the total number of genes with integrated provirus). This data suggests that (for example) integration into the NOSIP gene is more likely to generate a latent provirus than integration into the PACS1 gene (*p* = 2.3×10^-4^). The distance to the nearest host gene transcription start site (TSS) has been previously shown to be correlated with expression of the integrated provirus ^37^. For proviruses found within genes, we found that the distance to the TSS was smaller for latent than active proviruses, irrespective of orientation relative to host transcription (Fig. S12a-c). Proviral orientation relative to host gene transcription did not strongly influence whether proviruses were active or latent (Fig. S12d-f). Integration of proviruses into SPAD versus LAD regions did not lead to significant differences in their propensity to establish latency (Fig. S13a). Similarly, there was no significant difference in the proportion of active and latent proviruses when integration sites in distinct nuclear sub-compartments, defined using chromatin conformation capture (Hi-C) sequencing data ^63^, were compared (Fig. S13b). However, mapping of integration sites with respect to genomic repeat elements showed that latent proviruses were more frequently integrated into centromeric satellite regions (3.3% of repeat sequence integration sites) as compared to active proviruses (0.31% of repeat sequence integration sites) (Fig. 4g). This difference was also evident when integration sites in expanded clones or non-expanded clones were considered separately. In expanded clones, 13.6% of integration sites with latent proviruses and only 2.8% of integration sites with active proviruses were in centromeric satellite regions (Fig. S13c-d). Overall, the anatomical nuclear region did not detectably impact the propensity of HIV-1 proviruses to establish latency. However, discernible differences in the local sequence context in which transcriptionally active and latent proviruses were found show that the HIV-1 integration site influences the probability that a given provirus will establish latency.

### Repressive epigenetic modifications are enriched in proximity to latent proviruses

Epigenetic modifications in the vicinity of the integration site might constitute the underlying mechanism by which local sequence context affects proviral transcription. After identifying HIV-1 integration sites associated with active and latent proviruses, we examined the levels of histone modifications proximal to each integration site (up to 100 kb upstream or downstream of the integration site) using a previously published ChIP-seq dataset for primary CD4^+^ T cells^64^. Consistent with the fact that most integration sites are in genes, HIV-1 proviruses recovered from engrafted cells were preferentially found in the vicinity of genomic regions enriched for histone modifications typically found within gene bodies (H3K36me3) and enhancers associated with active transcription (H3K4me1 and to a lesser extent H3K27Ac and H3K4me3) (Fig. 5a and S14a). The repressive histone mark H3K27me3 was relatively depleted in the vicinity of integration sites, while another repressive histone mark H3K9me3 appeared unchanged in proximity to integration sites (Fig. 5a). The level of CpG DNA methylation was reduced proximal to integration sites overall but was marginally higher in the proximity of latent proviruses than active proviruses (Fig. 5a). These findings are in agreement with previous studies^53,65^, and the notion that HIV-1 integration is favored in transcriptionally active regions of the genome. However, there were not striking differences in the histone modifications associated with active versus latent proviruses when histone modifications were considered individually and proviral populations were considered as a whole.

**Figure 5.**
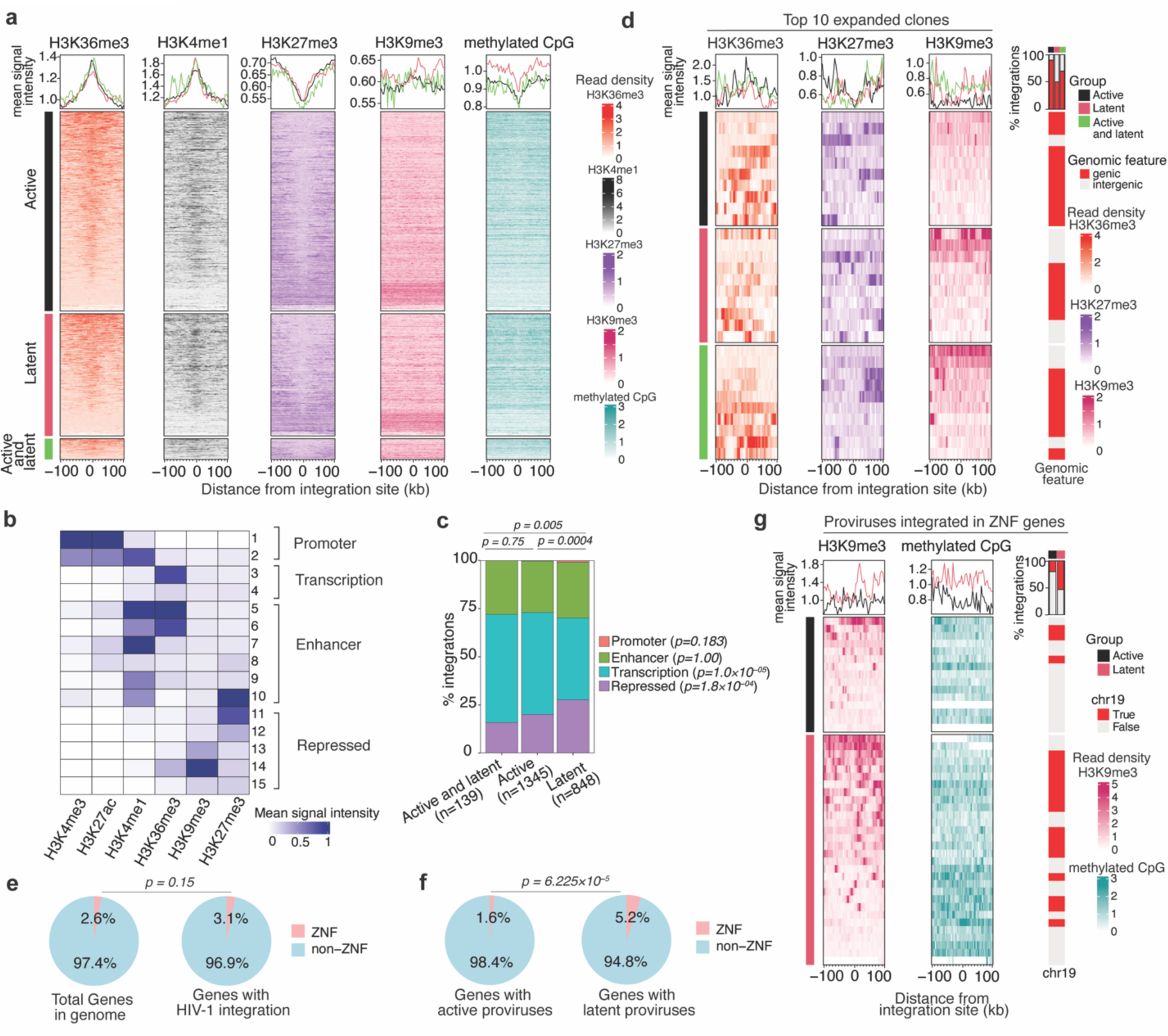
Distinct epigenetic signatures in the proximity of active and latent proviruses. **(a)** Heat map showing levels histone marks and methylated CpG found up to 100kb upstream and downstream of the integration sites of active, latent, or both (active and latent) proviruses. The line plot above the heat map shows the mean signal across all integration sites found within the active (black), latent (pink), and both (green) categories. The heatmaps for each group are arranged from top to bottom according to the highest to lowest mean signal intensity for H3K36me3 (orange) and this order is maintained for other histone marks, H3K4me1 (black), H3K27me3 (purple), H3K9me3 (pink), and methylated CpG (cyan), respectively. **(b)** Heatmap showing the mean signal intensity of various histone marks in the 15 chromatin states predicted by the ChromHMM model. **(c)** Stack bar plot showing the distribution of proviruses in distinct chromatin state groups. Global group-level statistical comparisons are shown on top of the plot using the chi-square test, and per-state level comparison between active and latent proviruses is given in in the legend using a two-proportion z-test. **(d)** The levels of H3K36me3 (orange), H3K27me3 (purple), and H3K9me3 (pink) histone marks in the proximity of the integration sites for active, latent, active and latent proviruses found in the top 10 expanded clones arranged in order arranged from top to bottom according to mean H3K9me3 signal. Bars to the right depict whether each proviruses within a group was integrated into a genic or intergenic region. **(e)** Pie charts shows the percentage of ZNF genes in the genome and the fraction of ZNF and non ZNF genes that have HIV-1 integration. **(f)** Proportion of ZNF and non-ZNF genes harboring either active or latent proviruses. The p-values were determined by Fisher’s exact test**. (g)** Histone modification levels proximal to active, latent, or active and latent proviruses found within ZNF genes.

Proviral integration sites are flanked by chromatin bearing multiple epigenetic modifications, whose effects on proviral expression may be exerted combinatorially. To integrate these signals and classify genomic regions into distinct chromatin states that account for combinatorial effects, we applied the generative machine-learning model ChromHMM^66^. The model uses epigenomic ChIP-seq datasets and enabled us to annotate the genome in CD4^+^ T-cells into 15 distinct chromatin states based on the signal intensity of 6 histone marks (Fig. 5b). These states can be grouped into four functional categories typically associated with promoter, enhancer, transcribed and repressed sequences. The occurrence of active and latent proviral integration sites within these chromatin-state defined categories showed that latent proviruses were depleted relative to active proviruses in genomic regions associated with host gene transcription marked by H3K36me3 (states 3 and 4) (Fig. 5b-c). Conversely, latent proviruses were found to be enriched within chromatin states associated with heterochromatin that is transcriptionally repressed, marked by H3K27me3 and H3K9me3 (state 11-15) (Fig. 5b-c).

We next compared the epigenetic features surrounding the integration sites of the 10 most expanded clones in the active, latent, and mixed (active and latent) categories. The integration sites were chosen based purely based the number of times they were encountered in the sorted categories; by restricting analyses to a population that was sampled most intensively, the transcriptional phenotype associated with each integration site could be ascribed with the greatest confidence. For the active, latent, and mixed (active and latent) categories, 9/10, 5/10, and 7/10 of the integration sites were in genes (Fig. 5d). Several of the latent and the mixed integration sites in this population were associated with intergenic genomic regions with abundant H3K9me3, but none of the 10 active proviruses demonstrated this property (Fig. 5d and S14b). Other epigenetic modifications did not differ noticeably among the 10 most expanded clones three categories.

### Properties of ZNF genes favor the occurrence of latent HIV-1 proviruses

Genes encoding ZNF proteins have recently been reported to harbor intact proviruses in individuals on long-term ART^23,29^. Importantly, ZNF genes are exceptionally abundant in the human genome, representing 2.6% of all genes. HIV-1 integration *per se* was not biased to ZNF genes and the large numbers of integration sites found therein (3.1% of all integrations) was approximately as expected based on the number of ZNF genes in the human genome (p = 0.15) (Fig. 5e). However, integration into a ZNF gene was more likely to lead to proviral latency than was integration in other genes. Among proviruses integrated within genes, ZNF genes harbored a higher fraction that were latent. Indeed, 5.2% of latent proviruses in genes but only 1.6% of active proviruses in genes were integrated in ZNF genes (Fig. 5f).

Notably, the expression of ZNF genes in general was lower than that of typical non-ZNF genes in CD4^+^ T-cells (Fig. S15a). and lower that that of genes that were typically favored for integration (Fig. S15 b-c). This conclusion held for ZNF genes that harbored active or latent proviruses (Fig. S15b-c). There was no statistically significant difference in the expression levels or the levels of promoter and gene body-associated histone marks for ZNF genes that had active versus latent proviruses (Fig. S15d-e). However, ZNF genes with latent proviruses had higher levels of integration site-proximal H3K9me3 compared to those of active proviruses (Fig. 5g). The levels of methylated CpG DNA were higher proximal to latent proviruses with integration sites in ZNF genes as compared to active proviruses (Fig. 5g) and this property was particularly prominent for latent proviruses in ZNF genes that did not exhibit proximal H3K9me3 enrichment. Overall, the majority of latent proviruses in ZNF genes were associated with proximally elevated levels of either H3K9me3 or methylated CpG DNA (Fig. 5g). These data clearly suggest that the type of epigenetic modifications found near the site of HIV-1 integration in the human genome affects the transcriptional fate of the integrated provirus and drives proviral latency. ZNF genes preferentially harbor latent proviruses due to their large numbers, comparatively low levels of expression, and enhanced levels of repressive H3K9me3 and CpG epigenetic modifications.

## Discussion

Here, we describe a model (HIVLM) to study the impact of HIV-1 integration site on proviral latency. In this model, memory CD4^+^ T-cells, each carrying a single transcriptionally active provirus, are engrafted into mice. The integrated proviruses in this model are fully competent to report HIV-1 gene expression but do not further replicate – no new viral particles or proviruses are generated after engraftment. Non-essential genes that may be cytocidal were removed from the viral genome, and the only viral gene products expressed are those that regulate gene expression (Tat and Rev). Moreover, the mice into cannot mount an immune response. Therefore, it is expected that little, if any, negative selective pressure that would favor transcriptional silencing is applied to the proviral population. Rather, the proviral population is unencumbered and can exert effects on the expansion of cell clones, and adopt transcriptionally active or latent states, based purely on the physiology of the cell clone that carries it, and the site in the host genome at which it is integrated.

Clonal expansion is an inherent phenomenon associated with memory T cell survival *in vivo*^67^ and provirus-containing cells clonally expand to varying degrees after engraftment. Notably though, a central finding of our study was that the integration site in cells harvested from mouse grafts mirrored the integration site profile during initial infection of CD4^+^ T cells prior to engraftment. Gene features that are key determinants of integration frequency *in vitro* including size, expression level, and nuclear localization, are sufficient to explain the occurrence of ‘recurrent integration genes’ such as NPLOC4, PACS1, NOSIP, and NFATC3 in grafted cells, which match those observed in human studies^11,23,51^. Conversely, TCR analysis revealed that some common T cell clones are favored for survival and expansion both HIV-1-infected and uninfected grafts. Overall, our results suggest that TCRs are drivers of T cell clonal expansion in our model and potentially in humans, while the HIV-1 provirus acts as a passenger during cell proliferation, in most cases.

The diversity of integration sites harboring latent or active proviruses sites in post-engraftment cells indicates that many locations in the human genome can sustain proviruses in either an active or a latent state. Nevertheless, there was a clear enrichment of transcriptionally latent proviruses in genomic regions that are repressed prior to integration such as intergenic and centromeric satellite regions. This observation is consistent with studies in which intact proviruses present in reservoir cells ART-treated human were found to be enriched in transcriptionally repressed regions in the genome^29,68,69^. The integration site profile in the natural reservoir after long-term ART presumably reflects the impact of immune selection pressure on a non-replicating viral population, favoring the survival of proviruses that are poorly expressed^29^. In the HIVLM model, where immune selection pressure is absent, unselected engrafted cells exhibit an integration profile indistinguishable from that of *in vitro* infection. Immune selection is mimicked experimentally in the HIVLM model, by the artificially application of selection pressure using flow cytometry or magnet-based sorting of harvested SBP^-^/GFP^-^ versus SBP^+^/GFP^+^ cells. That the impacts of immune selection over years in humans can apparently be mimicked in our HIVLM model after a single reporter gene-based simple sort, 2 months post engraftment is notable, and our approach enables many proviruses, with a great variety of integration sites to be evaluated.

A limitation of our study is that we did not directly measured epigenetic modifications at the provirus promoter or in the linear proximity of provirus integration site in the infected cell clones, but rather relied on the distribution of histone markers measured elsewhere in primary CD4 T^+^ cells^64^. Nevertheless, our data identifies host epigenetic signatures near provirus integration sites that predict latency. In some cases, integration associated epigenetic signatures (such as those found within genes, including elevated H3K36Me3, H3K4Me1, and reduced H3K27Me3) disfavor latency. Conversely, latency associated signatures include histone modifications, such as H3K9Me3 as well as CpG methylation, that are generally found outside genes, and typically encountered in heterochromatin.

The underlying drivers of the phenomenon, that latent intact proviruses are frequently found in ZNF genes^23,28^, is suggested by our data. ZNF genes are a particularly abundant gene class (3.6% of all genes), and our data shows that integration in ZNF genes predisposed integrated proviruses to acquire latency in the HIVLM model. Notably ZNF genes exhibit reduced expression in CD4+T-cells than is typical of human genes of a whole. Moreover, genomic loci in the vicinity of the subset of ZNF genes that were found to harbor latent proviruses in our experiments exhibited signatures of an epigenetically repressed states. These states included with a high level of H3K9me3 mark, or higher than typical levels of methylated CpG. We note that about 40% of the >700 ZNF genes in the human genome reside on a single human chromosome (chromosome 19), and while chromosome 19 is exceptionally gene dense, ∼20% of all genes on chromosome 19 are ZNF genes. Chromosome 19 is also relatively G+C rich and is enriched in CpG dinucleotides and islands^70^. Chromosome 19 was also strikingly enriched in latent proviruses in the HIV-LM model. We posit that ZNF genes represent an abundant gene class, residing in a are of genome where a confluence of factors favoring HIV-1 integration such as high gene density gene density and those favoring latency (low gene expression and a repressed chromatin state) are balanced in such a way that the probability of generating latent proviruses approaches a maximum.

Overall, the approaches described herein provide a model system in which epigenomic mechanisms involved in latency can be studied and HIV-1 cure strategies employing approaches to reverse latency in reservoir cells can be evaluated.

## Material and methods

### Proviral construct V1/SBP-GFP

A proviral plasmid pV1/SBP-GFP was generated based on a minimal viral genome (V1)^71^, engineered from HIV-1 NL43 harboring large deletions or inactivating mutations in *gag*, *pol*, *vif*, *vpu*, and *env*, and in which *nef* was replaced by GFP. To construct the V1/SBP-GFP plasmid, an DNA fragment encoding a streptavidin binding peptide appended to the extracellular domain of the low affinity nerve growth factor receptor and a P2A site, flanked by NotI restriction sites was synthesized, PCR-amplified, and cloned into the NotI-linearized pV1 plasmid.

### Cell lines

293T cells (ATCC, #CRL-3216; RRID: CVCL_0063) and MT4-LTR-GFP cells^72^ were cultured at 37 °C and 5% CO2 in DMEM (293T) or RPMI (MT4-LTR-GFP) supplemented with 10 % FCS and Gentamycin. All cell lines used in this study were monitored regularly to ensure the absence of retroviral contamination. Cells were also monitored regularly for mycoplasma contamination. Cells used here were contamination-free.

### Virus production and characterization

To generate V1/SBP-P2A-GFP-derived viral stocks, 7.5×10^7^ 293T cells (ATCC, #CRL-3216; RRID: CVCL_0063) were seeded per 875cm^2^ 5-layer flask the day before transfection. Cells were transfected with 75 µg pV1/SBP-P2A-GFP, 75 µg pCRV1/NLGagPol, and 15 µg pVSV-G^73^ using polyethyleneimine. 12 h post-transfection, medium was replaced with fresh growth medium. Virus-containing supernatants were harvested and filtered 48 h post-transfection. Virus was pelleted by ultracentrifugation though a 20% sucrose cushion, resuspended in PBS with 10% BSA to achieve 70-80-fold concentrated virus stocks, aliquoted, and stored at -80 °C. Infectious titers were determined by titration on MT4-LTR-GFP cells^72^ and assessment of %GFP-positive cells by flow cytometry 48 hrs after infection.

### Isolation and activation of memory CD4^+^ T cells

A small-volume blood draw (50 ml whole blood) and Leukapheresis were performed at the Rockefeller University Hospital from a healthy adult participant who provided written informed consent before participation. Samples were processed within 4 hrs of collection. PBMCs were isolated from 50 ml whole blood (test infection to determine optimal participant-specific MOI) or 100 ml Leukopack (for preparation of cells for engraftment) by density gradient centrifugation using lymphocyte separation medium (Corning). Memory CD4^+^ T cells were isolated from PBMCs using the EasySep Human Memory CD4^+^ T Cell Enrichment Kit (StemCell, #19157). A subset of cells was used for purity assessment of memory CD4^+^ T cells (CD4^+^CD45RA^-^CD45RO^+^) by flow cytometry using the following fluorochrome-conjugated antibodies: anti-human CD4 Antibody, clone OKT4 (BioLegend, #317408), anti-human CD45RA Antibody, clone HI100 (BD, # 550855), and anti-human CD45RO Antibody, clone UCHL1 (BD, #555493). Approximately 25 % of retrieved memory CD4^+^ T cells from Leukopack were frozen at a density of 1×10^7^ cells/ml in 100% fetal bovine serum and stored at -150 °C. The remaining memory CD4^+^ T cells were cultured in regular RPMI growth medium supplemented with IL-2 (RPMI supplemented with 10% fetal bovine serum, gentamycin, 2 ng/ml (20 U/ml) human recombinant IL-2 (PeproTech, #200-02)) and activated with 25 µl Dynabeads Human T-Activator CD3/CD28 (Thermo Scientific) per 1×10^6^ cells. Dynabeads were removed after 48 hrs, and cells were prepared for infection. Activation was assessed by flow cytometry using anti-human CD69 antibody (BD, #555533).

### Infection of memory CD4^+^ T cells

Following removal of CD3/CD28 T cell activator beads, 1.3×10^9^ memory CD4^+^ T cells were infected using 2×10^7^ IU (based on titration of MT4-LTR-GFP cells, equivalent to an approximate participant-specific MOI of 0.3) V1/SBP-GFP virus in the presence of 8 µg/ml polybrene. For this, the virus was diluted in 50 ml RPMI growth medium supplemented with IL-2, and polybrene was added at approximately. 9x concentration. The virus/polybrene mixture was added to the cells, which were adjusted for a final cell density of 3×10^6^ cells/ml in fresh RPMI growth medium supplemented with human recombinant IL-2. After 12 hrs, the cell density was decreased to 1×10^6^ cells/ml using fresh RPMI growth medium supplemented with human recombinant IL-2. Infection efficiency was assessed 48 hours after infection by flow cytometry (GFP).

### Antibody-free magnetic isolation of HIV-reporter-expressing memory CD4^+^ T cells

Cells were washed once with PBS and once with PBS + 2mM EDTA + 0.2 % BSA. Cells were resuspended at a density of 1×10^7^ cells/ml in pre-warmed DMEM + 2mM EDTA + 0.2 % BSA and incubated with pre-washed Dynabeads Biotin Binder (Thermo Fisher Scientific, #11047) at a GFP+ cell: bead ratio of 1:20. Following 30 min incubation shaking at 37 °C, unbound (negative) cells were removed and bead-bound (positive) cells were washed twice on a magnet using pre-warmed DMEM + 2mM EDTA + 0.2 % BSA. For removal of the beads from the isolated positive cells, bead-bound cells were incubated in pre-warmed RPMI growth medium supplemented with 2mM biotin and incubated for 15 min at 37 °C on a shaker. Beads were removed using a magnet and once again incubated with biotin-containing medium. Supernatants containing released cells were pooled, counted, and the cell density was adjusted to 5×10^5^ cells/ml in RPMI growth medium supplemented with IL-2. The efficiency of antibody-free magnetic cell sorting was confirmed by analysis of input, negative, and positive cells using flow cytometry (GFP).

### Preparation of cells for engraftment

At 48 hours after antibody-free magnetic isolation of V1/SBP-GFP-expressing cells, the purity of cells was confirmed by flow cytometry, and cells were frozen for future engraftment at a density of 1×10^7^ cells/ml, 1 ml/tube in fetal bovine serum with 10% DMSO. Cells were stored at -150 °C until engraftment.

The day before engraftment, V1/SBP-GFP-expressing memory CD4^+^ T cells frozen in fetal bovine serum were thawed and quickly resuspended in 7-fold excess volume of pre-warmed RPMI growth medium and centrifuged at 300g for 10 min. Cells were adjusted to 1.6E6 cells/ml in RPMI growth medium supplemented with 50 U/ml IL-2 and cultured for 14 hrs. Before engraftment, cells were washed once with PBS and resuspended in HBS at 6E7 cells/ml. 100 µl of this cell suspension (6E6 cells) was used per mouse for engraftment.

### Engraftment of NSG mice with V1/SBP-GFP-expressing human memory CD4^+^ T cells

Irradiated (50 rad) 6-week-old NSG mice (NOD.Cg-Prkdc^scid Il2rg^tm1Wjl/SzJ from The Jackson Laboratory) were engrafted with 6×10^6^ V1/SBP-P2A-GFP-expressing memory CD4^+^ T cells resuspended in 100 µl HBS by tail vein injection. Mice were periodically bled to assess human immune cell reconstitution and V1/SBP-GFP expression. Approximately 100 µl of peripheral blood was collected in EDTA-coated tubes. Cell pellets were stained for flow cytometric analysis. Mice were cared for by trained research animal technicians and received daily wellness checks. Procedures were performed in accordance with protocols approved by the Weill Cornell Medical College Institutional Animal and Use Committee and Rockefeller University CBC facility (Protocol # 24016-H). At 63 days after engraftment, mice were sacrificed, and blood, bone marrow, lung, and spleen were isolated. Cells were immunophenotyped for human CD45, CD3, CD8, and CD4 expression, as well as V1/SBP-GFP expression. Remaining harvested cells were frozen at -80 °C until DNA extraction.

### Flow cytometric analysis of human cells engrafted in NSG mice

Pre-engraftment samples were analyzed for purity using a panel of the following human-specific antibodies: FITC anti-CD4 (OKT4), PE anti-CD45RO (UCHL1), and APC anti-CD45RA (HI100). T cell activation was analyzed using APC anti-CD69, and V1/SBP-GFP expression was analyzed by detection of GFP. For engraftment samples, the mice’s peripheral blood was collected into EDTA-coated tubes from the mice by submandibular bleed every 2 weeks to assess human T cell reconstitution. Blood was centrifuged at 2000g for 5 mins to separate plasma and cell pellets. Cells were immediately stained for flow cytometric analysis. Samples were stained with human-specific antibodies for the analysis of cell surface markers: CD4 BV421, CD8 BV605, CD3 BV711, CD45 PerCp-Cy5.5, CD45RO-PE, CD69 PE-eFluor 610, CCR7 PE-Cy-7, CD27 APC, CD25 APC-eFluor780. Fluorescent counting calibration beads were added to each sample to determine absolute cell counts. For post-engraftment spleen sample analysis and sorting, a similar antibody panel was used in combination with Live/Dead Aqua (Invitrogen) dead stain as a cell viability marker. Stained samples were run on Attune NxT cytometer analyzer (Thermo Fisher Scientific), sorted on BD FACS Aria III (BD Biosciences), and data were analyzed by FlowJo version 10 software (Treestar, Ashland, OR).

### Intact Proviral DNA Assay (IPDA)

Genomic DNA was isolated from blood, bone marrow, lung, and spleen using the Zymo Quick-DNA miniprep PLUS kit according to the manufacturer’s instructions. Intact HIV-1 copies/million CD4^+^ T cells or per ng DNA were determined by digital droplet PCR (ddPCR) using the Intact Proviral DNA Assay (IPDA)^49^, where HIV-1 and human RPP30 reactions were performed independently in parallel, and copies were normalized to the amount of input DNA. In each ddPCR reaction, an average of 15 ng of genomic DNA was combined with ddPCR supermix for probes (no dUTP, BioRad), primers (final concentration 900 nM, Integrated DNA technologies), probes (final concentration 250 nM, Thermo Fisher Scientific), and nuclease-free water. Primer and probe sequences (5′–>3′) were as follows:

RPP30 Forward Primer- GATTTGGACCTGCGAGCG,

RPP30 Probe- VIC-CTGACCTGAAGGCTCT- MGBNFQ,

RPP30 Reverse Primer- GCGGCTGTCTCCACAAGT;

HIV-1 Ψ Forward Primer- CAGGACTCGGCTTGCTGAAG,

HIV-1 Ψ Probe- FAM- TTTTGGCGTACTCACCAGT- MGBNFQ,

HIV-1 Ψ Reverse Primer- GCACCCATCTCTCTCCTTCTAGC;

HIV-1 env Forward Primer- AGTGGTGCAGAGAGAAAAAAGAGC,

HIV-1 env Probe- VIC-CCTTGGGTTCTTGGGA- MGBNFQ,

HIV-1 env Reverse Primer- GTCTGGCCTGTACCGTCAGC.

Droplets were prepared using a QX200 Droplet Generator (BioRad) and cycled at 95 °C for 10 min; 45 cycles of (94 °C for 30 sec, 59 °C for 1 min) and 98 °C for 10 min. Droplets were analyzed on a QX200 Droplet Reader (BioRad) using QuantaSoft software (BioRad). Four technical replicates were run for each sample.

### Ligation-mediated PCR-based HIV-1 integration site analysis

A detailed procedure for intergrtaion site determination using PRISM-seq will be published elsewhere (manuscript submitted). Briefly, extracted DNA was diluted according to ddPCR results, so that 50 provirus was present in each well of the 96-well plate. Subsequently, gDNA in each well was subjected to multiple displacement amplification (MDA) with phi29 polymerase (Qiagen REPLI-g Single Cell Kit #150345) at 30 °C for 4 h. DNA produced by whole-genome amplification was used as a template. Viral-host junction regions were amplified using a nested PCR strategy. 5LTR host junction was amplified with LSP1 (GCTTCAGCAAGCCGAGTCCTGCGTCGAG) and LSP2 (GCTCCTCTGGTTTCCCTTTCGCTTTCAA), and 3LTR host junction was amplified with LestraLTR1 (CTTAAGCCTCAATAAAGCTTGCCTTGAG) and LestraLTR2 (AGACCCTTTTAGTCAGTGTGGAAAATC) as forward primers in the 1^st^ and 2^nd^ PCR reaction in combination with reverse primers annealing with the ligated adaptor sequence, AP1 (GTAATACGACTCACTATAGGGC) and AP2 (ACTATAGGGCACGCGTGGT) respectively. The resulting PCR products were subjected to next-generation sequencing using Illumina MiSeq. MiSeq paired-end FASTQ files were demultiplexed; small reads (142 bp) were then aligned sequentially to HIV-1 reference genomes HXB2 then to the human reference genome GRCh38. Biocomputational identification of integration sites was performed using a house R-based pipeline (BulkIntSiteR). Briefly, chimeric reads containing both human and HIV-1 sequences were evaluated for mapping quality based on (i) HIV-1 coordinates mapping to the terminal nucleotides of the viral genome, (ii) absolute counts of chimeric reads, and (iii) depth of sequencing coverage in the host genome adjacent to the viral integration site. The final list of integration sites and corresponding chromosomal annotations was obtained using Ensembl (v86, www.ensembl.org), the UCSC Genome Browser (www.genome.ucsc.edu), and GENCODE (v39, www.gencodegenes.org). Repetitive genomic sequences harboring HIV-1 integration sites were identified using Repeat Masker (www.repeatmasker.org).

### TCR analysis of HIV-1 infected and uninfected cell populations

Briefly, 1ug gDNA samples from mouse spleen samples of HIV-1 infected and uninfected engrafts were outsourced to Adaptive Biotechnologies for TCR sequencing using the assay type hsTCRBv4b. The data obtained was analyzed using the ImmunoSeq analyzer and an in-house built-in pipeline. The clones with a read count of 1 were removed, and the CDR3 protein sequences were taken to create a unique list of TCR in each individual sample. The log count per million was used as a measure to indicate the degree of clonal expansion of each cell clone in an individual sample. The Pearson correlation analysis was performed to compare clones that expanded across the mice engrafts in HIV-1 infected and uninfected samples.

### Analysis of epigenetic modifications proximal to integrations sites

The HIV-1 integration site coordinates retrieved from cells harboring latent and active proviruses were used to generate 4kb binned chromosomal regions spanning 100kb upstream and downstream of the provirus. The ChiP-seq dataset was assessed from ENCODE database for H3K4me1, H3K4me3, H3K27ac, H3K36me3, H3K27me3, and H3K9me3 histone marks for primary CD4^+^ T cells using the following criteria: organ, blood; cell, leukocyte; biosample, GM12878 or CD4-positive, alpha-beta memory T cell; genome assembly, GRCh38; file type, “bigwig”; and output type, “fold change over control.” Only datasets based on two replicates are used. The histone modification signal for each epigenetic mark was represented as an average read signal for 100 kb upstream and downstream of the proviral integration coordinate using the UCSC bigWigAverageOverBed tool.

### Quantification and statistical analysis

Data are presented as pie charts, bar charts, scatter plots with individual values, or box and Whisker plots (indicating the median, minimum, maximum, and interquartile ranges). Differences were tested for statistical significance using Mann-Whitney U tests, Fisher’s exact tests, chi-square tests, and two-sided Kruskal-Wallis nonparametric tests, as appropriate. P < 0.05 was considered significant; false discovery rate (FDR) correction was performed using the Benjamini-Hochberg method. Statistical details can be found in the corresponding figure legends. Analyses were performed using GraphPad Prism and R software.

## Acknowledgements

We thank the Rockefeller University flow cytometry core, Comparative Biosciences Center (CBC), and Genomic Resource Center. We are grateful to Marina Caskey (The Rockefeller University) for organizing Leukapheresis at The Rockefeller University Hospital and Emily Mastrocola and Dennis Copertino for technical assistance. This work was supported by NIH grants to the Research Enterprise to Advance a Cure for HIV (REACH) 5UM1AI164565 and the Center for Structural Biology of HIV RNA U54 AI170660. This article is subject to HHMI’s Open Access to Publications policy. HHMI lab heads have previously granted a nonexclusive CC BY 4.0 license to the public and a sublicensable license to HHMI in their research articles. Pursuant to those licenses, the author-accepted manuscript of this article can be made freely available under a CC BY 4.0 license immediately upon publication.

---

**Figure S1.**
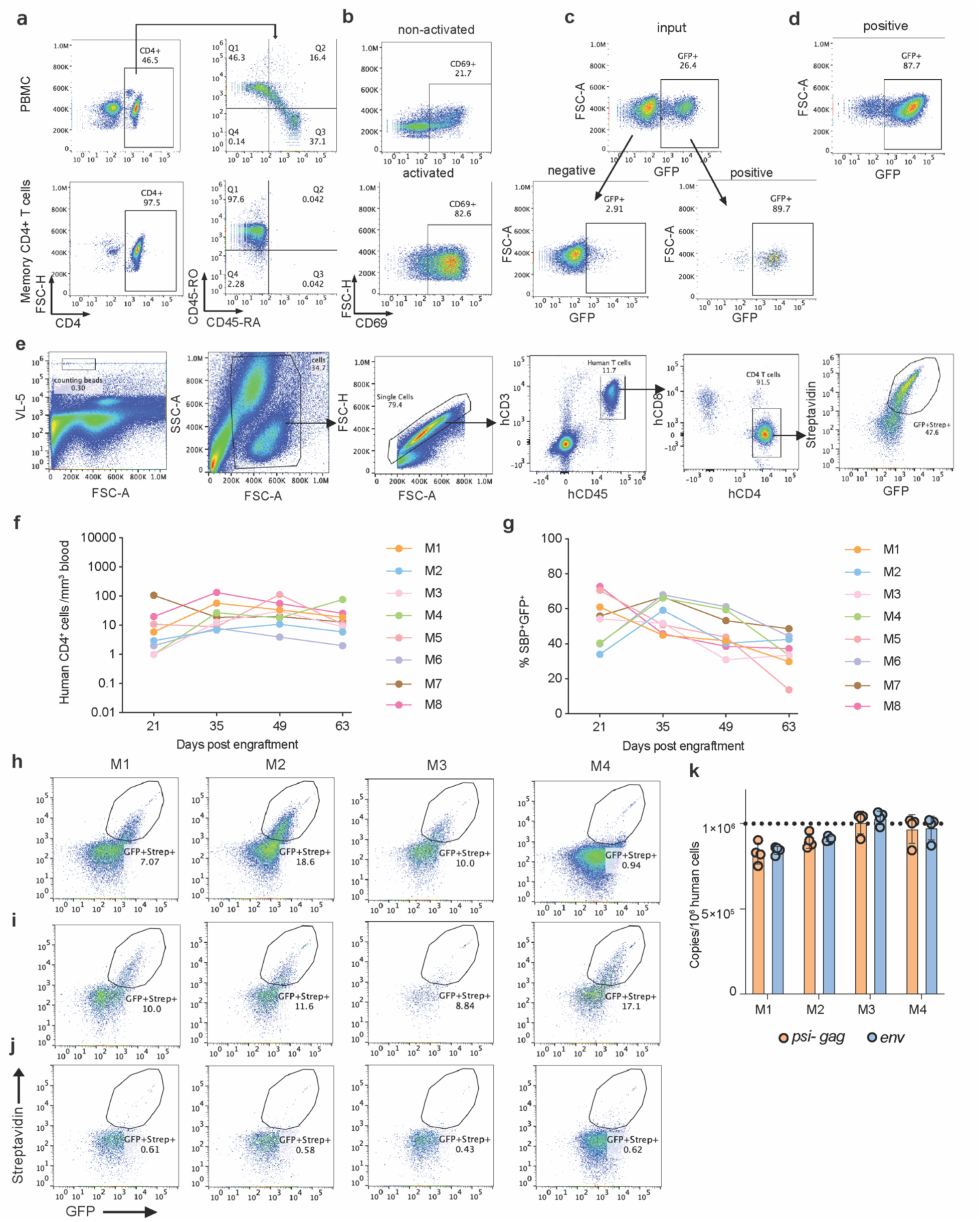
Infection, selection of reporter HIV-1-infected human memory CD4^+^ T cells, and engraftment of NSG mice. **(a)** Flow cytometry plot showing the isolation of human memory CD4^+^ T cells (lower panels) from peripheral blood mononuclear cells (PBMCs, upper panels) of an HIV-1-uninfected human donor. **(b)** Frequency of activated human memory CD4^+^ T cells (CD69^+^) 4 days after activation with anti-CD3/CD28 beads and 2 days after infection with V1/SBP-GFP virus. **(c)** Streptavidin magnetic bead-based cell isolation of V1/SBP-GFP-expressing memory CD4^+^ T cells. Bead-bound (positive) cells were separated from unbound (negative) cells 48 hours after infection with V1/SBP-GFP virus. **(d)** Percentage of V1/SBP-GFP positive cells 48 hours after sorting, immediately before freezing cells in aliquots for engraftment. **(e)** Flow cytometry gating strategy used for quantification of human CD4^+^ T cell counts and HIV-1 expression post engraftment. Counting beads for calibration of the quantification were gated based on FSC-A and VL-5 (detector), an Attune NxT detector excited by a 405nm laser with an emission filter at 710/50nm. The figure reports the percentage of transcriptionally active HIV-1-infected (SBP^+^GFP^+^) cells in the hCD3^+^/hCD45^+^/hCD4^+^ human cell compartment in a typical mouse peripheral blood sample. **(f, g)** Quantification of the numbers of human cells over ∼2 months following mouse engraftment; total number of human cells (**f**) and percentage of transcriptionally active HIV-1 infected (SBP^+^GFP^+^) human cells (**g**) over time in multiple animals. **(h-j)** Percentage of transcriptionally active HIV-1 infected (SBP^+^GFP^+^) human cells (gated on hCD3^+^/hCD45^+^/hCD4^+^ cells) in mouse spleen (**h**), lung (**i**), and bone marrow (**j**) of individual animals harvested 2 months after engraftment. **(k)** Quantification of *psi-gag* and *env* DNA copies per million human cells in spleen from each engrafted mouse.

**Figure S2.**
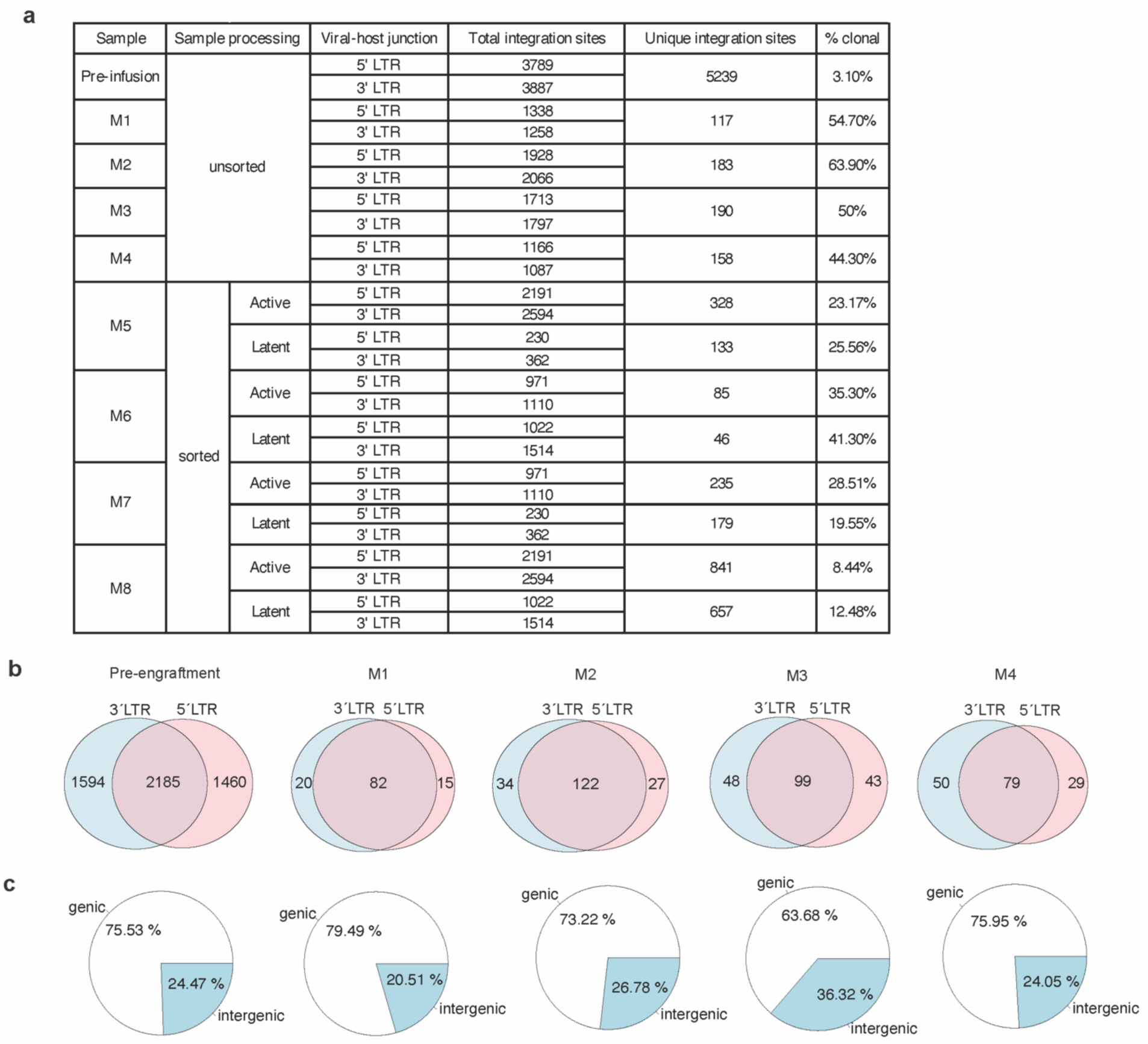
Identification of proviral integration sites in HIV-1 infected human memory CD4^+^ T cell populations pre- and post-engraftment. **(a**) Table showing the number of unique integration sites retrieved from human cells harvested from the spleen of individual, engrafted mice. Cells recovered from mouse spleens for animals M1-M4 were not sorted, while human cells recovered from mice M5-M8 were sorted based on SBP/GFP expression. The ‘percent clonal’ column reports the proportion of the total number of unique integration sites found within a sample that were detected in more than one well out of the 96 wells tested. **(b)** Comparison and overlap in the numbers of integration sites retrieved by either 5’LTR or 3’LTR viral-host junction amplification strategies for pre- and post-engraftment cell populations. **(c)** Distribution of integrated proviruses in genes and intergenic regions in pre- engraftment cells and post-engraftment cells from individual mice.

**Figure S3.**
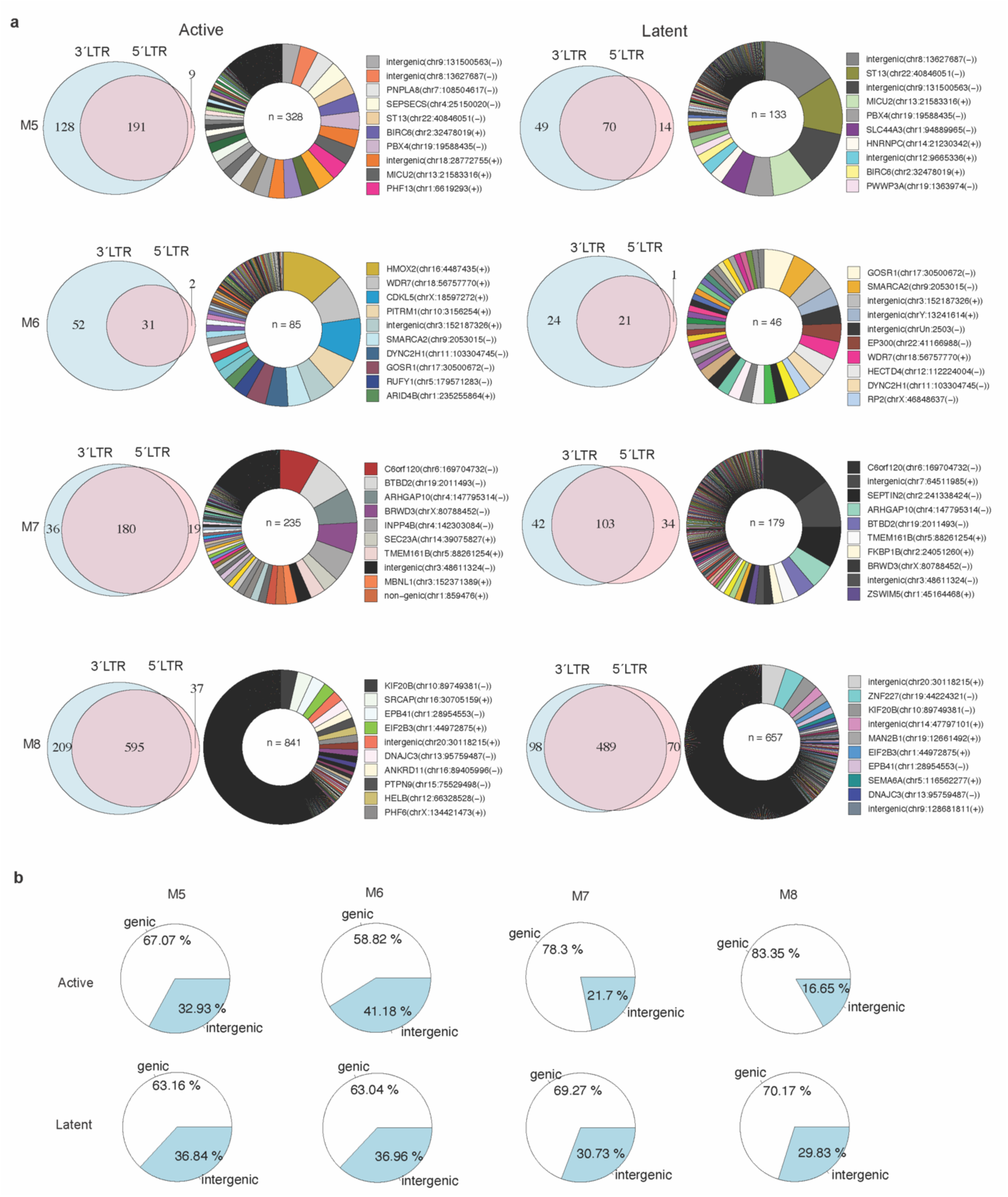
Integration sites identified in HIV-1 infected human memory CD4^+^ T-cells post-engraftment with active and latent proviruses. **(a)** Comparison of integration sites retrieved by 5’LTR and 3’LTR viral-host junction amplification. Circos plots showing the relative proportion of expansion of each clone in post-engraftment human cells from individual mouse spleens, sorted into populations with active and latent proviruses. Each clone is color-coded according to its proviral integration site. Each integration site is labelled as the name of the gene with integrated provirus or as intergenic if integration was outside of the gene, followed by the chromosome name, the genomic coordinate, and the host DNA strand (positive or negative). **(b)** Distribution of integrated proviruses in genes and intergenic regions in pre- and post-engraftment samples from individual mice.

**Figure S4.**
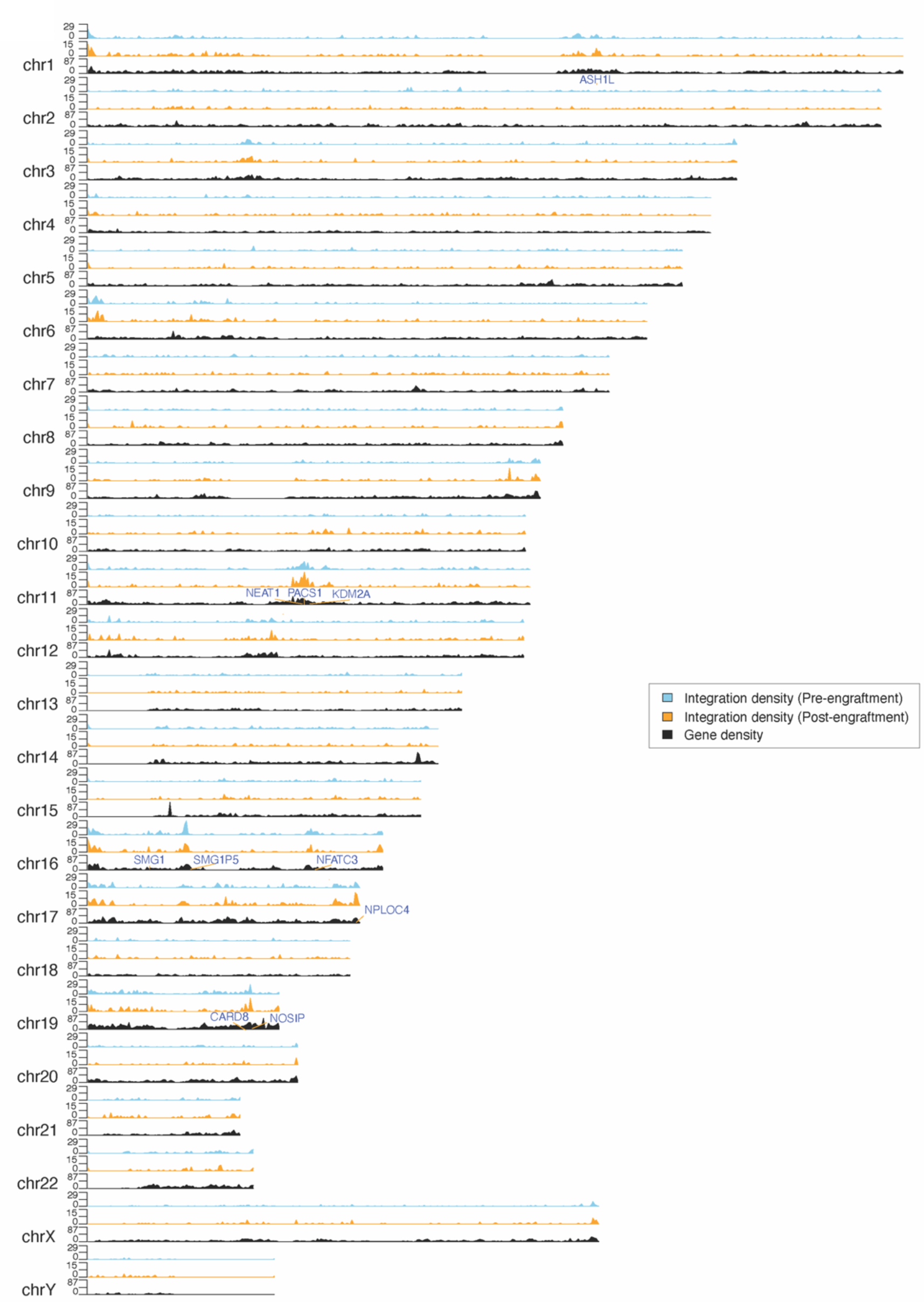
Gene density and integration frequency for human chromosomes and integration sites found in the pre- and post-engraftment cell populations. Genes with >5 unique integration sites post-engraftment are highlighted. The stacked y-axes represent the number of genes (0-87), the number of integrations in pre-(0-29), and post- (0-15) engraftment cell populations.

**Figure S5.**
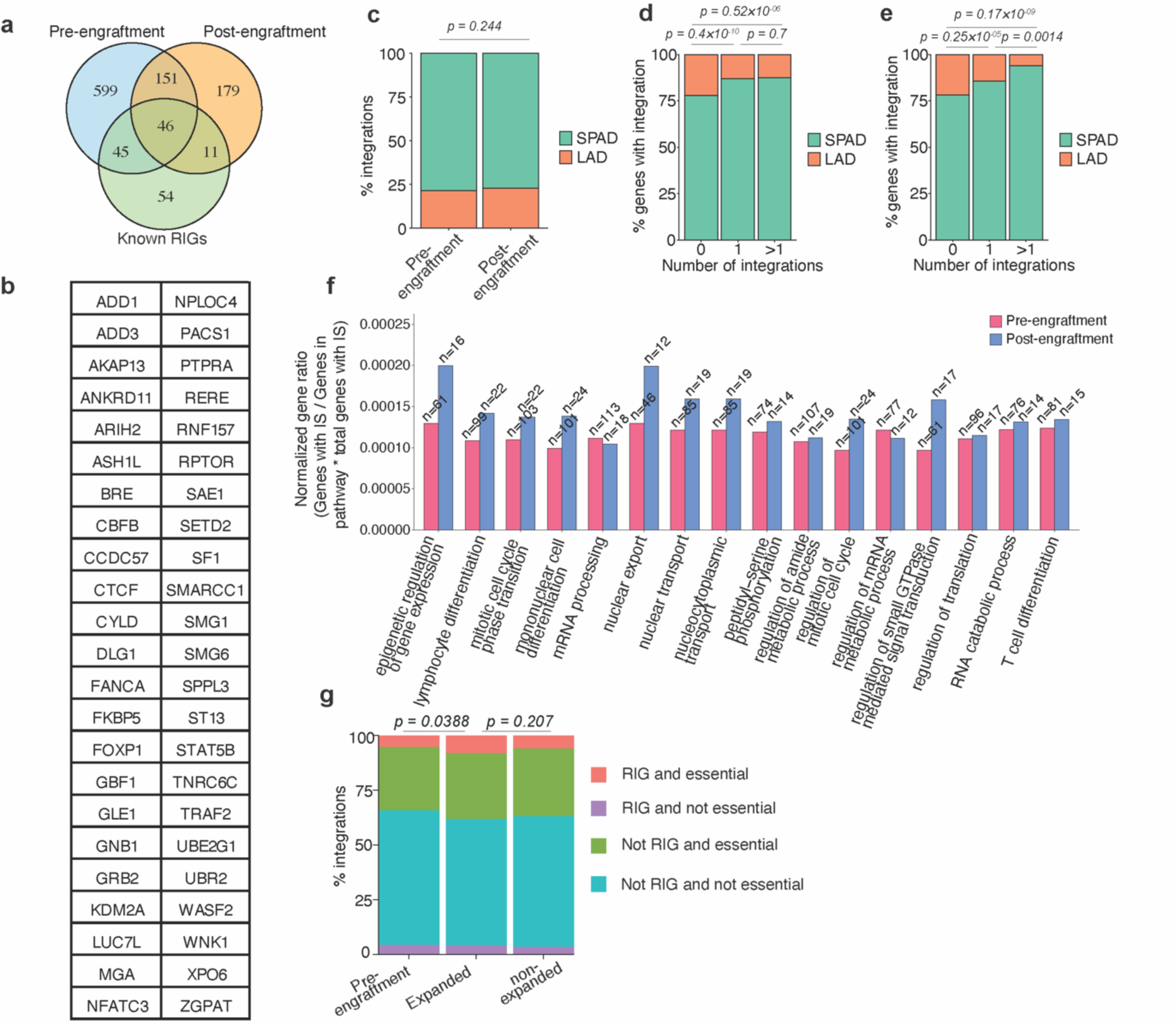
**(a)** Overlap of genes with >1 integration identified in the pre- and post- engraftment cell populations with genes designated RIGs in previous studies^51,55,56^. **(b)** List of 46 genes that overlap between the pre- /post-engraftment and the previously known RIGs. **(c)** Percentage of proviral integration in SPAD and LAD for integration sites obtained from pre- and post-engraftment populations. **(d, e)** Distribution in SPAD and LAD genomic regions for genes with 0, 1 or >1 integrations in pre- (d) and post-engraftment (e) samples. **(f)** Comparison of genes with HIV-1 integration sites in pre- and post-engraftment samples for association with various pathways as defined by gene ontology (GO) enrichment analysis. The top 15 enriched pathways based on the post-engraftment genic integration sites dataset were selected, and their respective enrichment in both pre- and post-engraftment samples compared. **(g)** Genic integration site distribution in genes that are RIG and essential, RIG and not essential, not RIG and essential, and not RIG and not essential, for pre-engraftment, as well as expanded, and non-expanded clones in post-engraftment datasets. P values determined by Pearson’s Chi-squared test.

**Figure S6.**
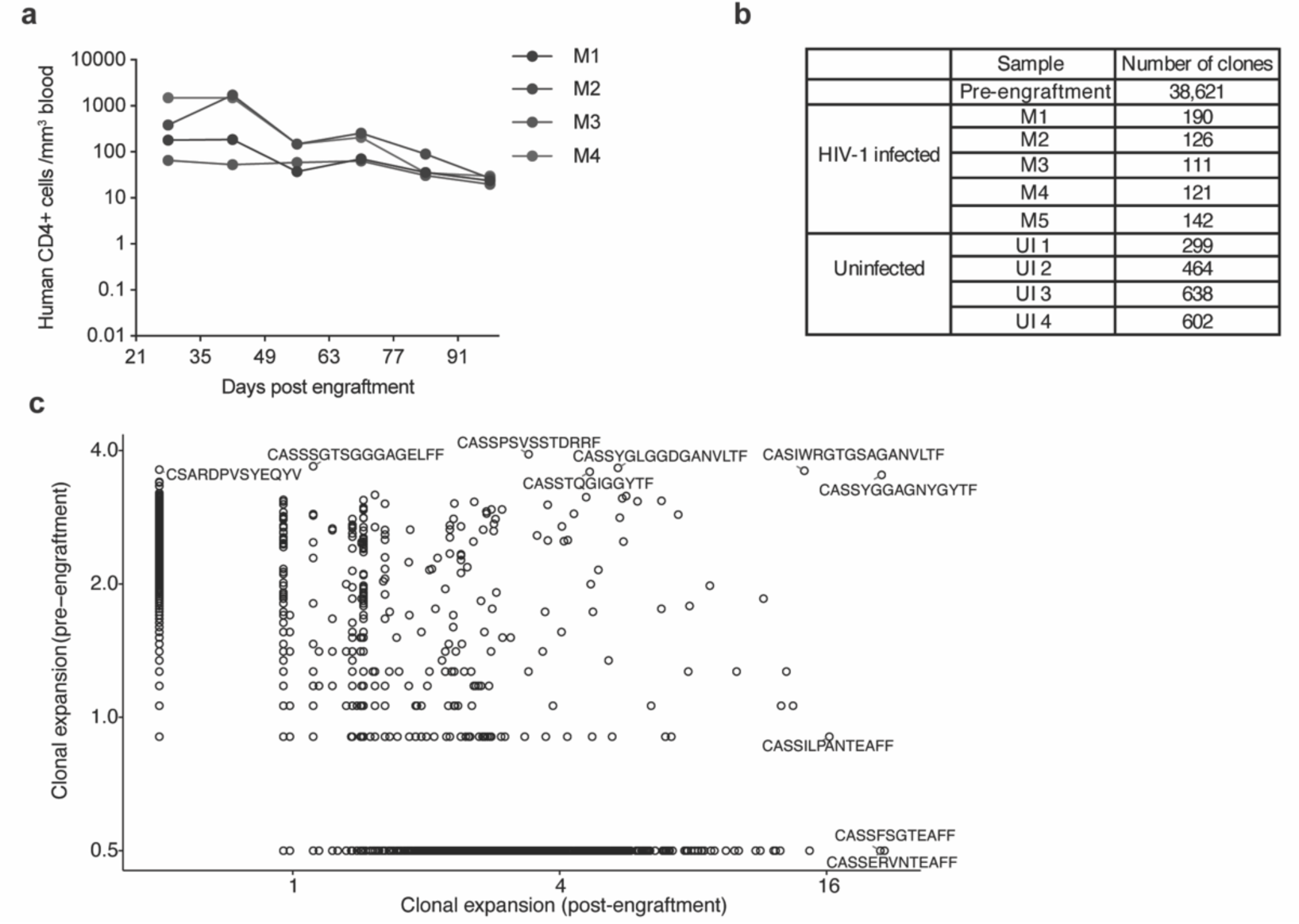
**(a)** Quantification of the number of human cells following engraftment of uninfected human memory CD4^+^ T cells over time in multiple animals. Each mouse was sacrificed at 98 days to measure the clonal expansion of individual T cell clones. **(b)** Number of T cell clones retrieved from pre-engraftment, HIV-1-infected, and uninfected post engraftment cell populations from individual mouse spleen samples. **(c)** Comparison of TCR sequence abundance in pre- and post-engraftment cell populations. The dot plot shows the total expansion of all clones retrieved from the uninfected and infected grafts. Each dot represents a T cell clone marked by its CDR3 TCR sequence. TCR sequence not detected in one of the two populations are assigned an arbitrary value of 0.5 for representation on a logarithmic scale.

**Figure S7.**
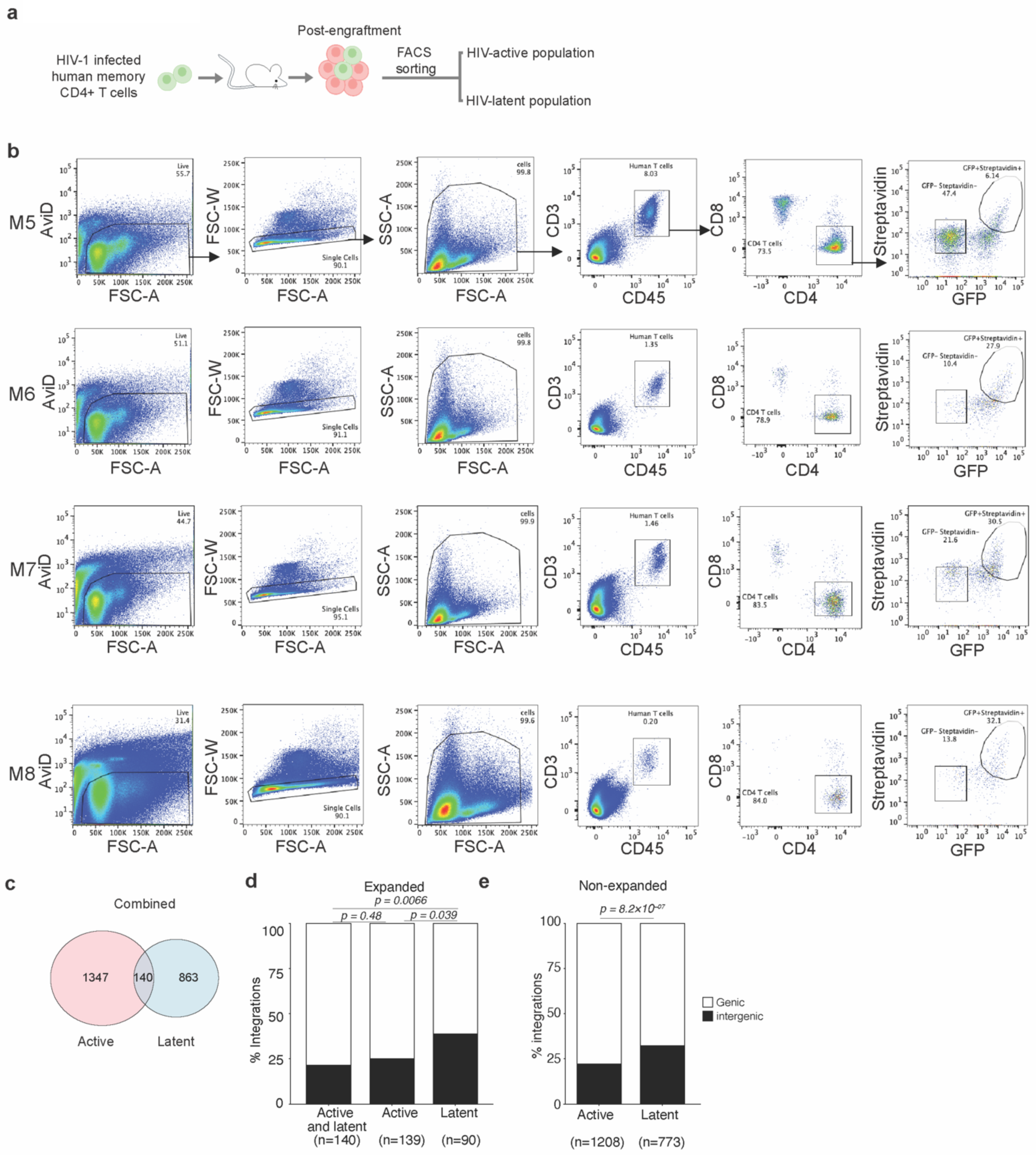
**(a)** Procedure for sorting of cell populations with active and latent HIV-1 proviruses post-engraftment. **(b)** Flow cytometry gating strategy to sort active (GFP^+^SBP+) and latent (GFP^-^SBP^-^) HIV-1-infected cell subpopulations post-engraftment. **(c)** Overlap analysis of integration sites found in sorted active and latent cell populations post-engraftment, pooled data from four mice. **(d, e)** Genic and intergenic distribution of integration sites obtained in active, latent, or active and latent proviruses for expanded and non-expanded cell clones, respectively. P-value denotes Fisher’s exact test.

**Figure S8.**
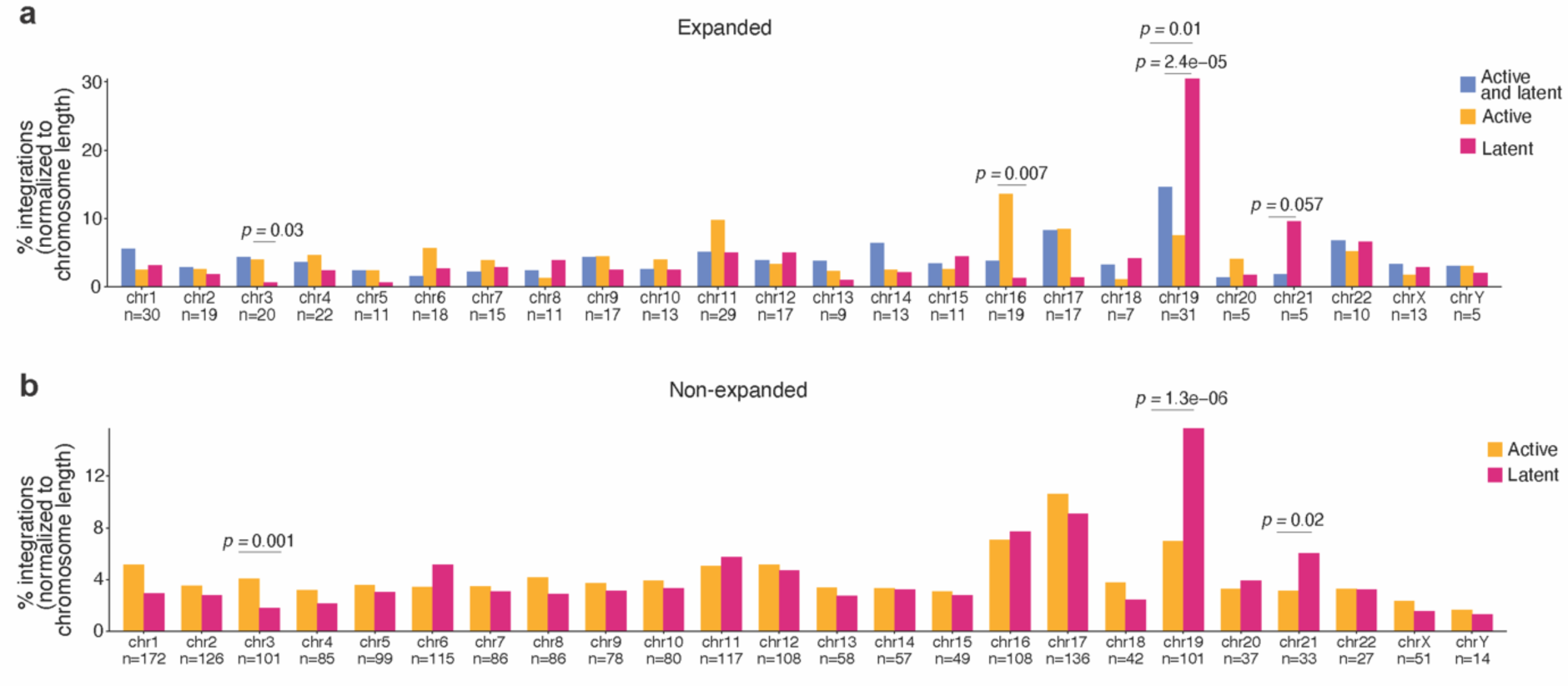
**(a** and **b)** Distribution of active, latent, or active and latent provirus integration sites found in expanded and non-expanded cell clones on human chromosomes. P-value denotes Fisher’s exact test.

**Figure S9.**
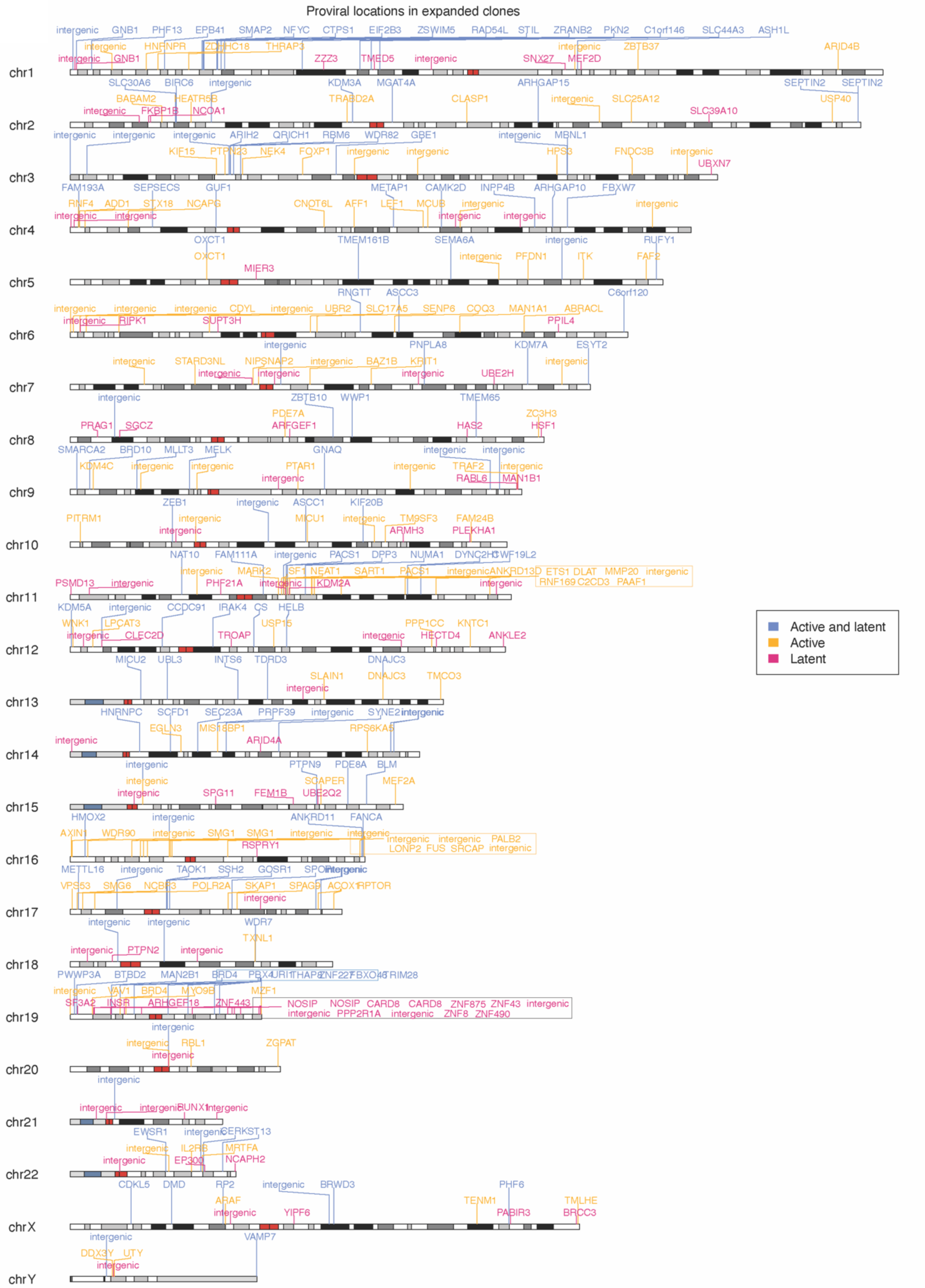
Karyoplot showing the chromosomal distribution of active, latent, or both active and latent provirus integration sites identified in expanded clones. Each ideogram represents a chromosome, and integration sites are mapped to their genomic coordinates. Proviruses are colored according to transcriptional status. Cytogenetic bands are shown based on Giemsa staining patterns following UCSC Genome Browser and ISCN conventions. Cytoband color key shows white (gneg), gene-rich GC-rich euchromatin; light to dark gray (gpos25–gpos100), increasing Giemsa-positive intensity indicating progressively AT-rich, gene-poor heterochromatin; red, centromeric regions; blue, secondary constrictions containing rRNA gene clusters; gray (gvar), variable heterochromatin.

**Figure S10.**
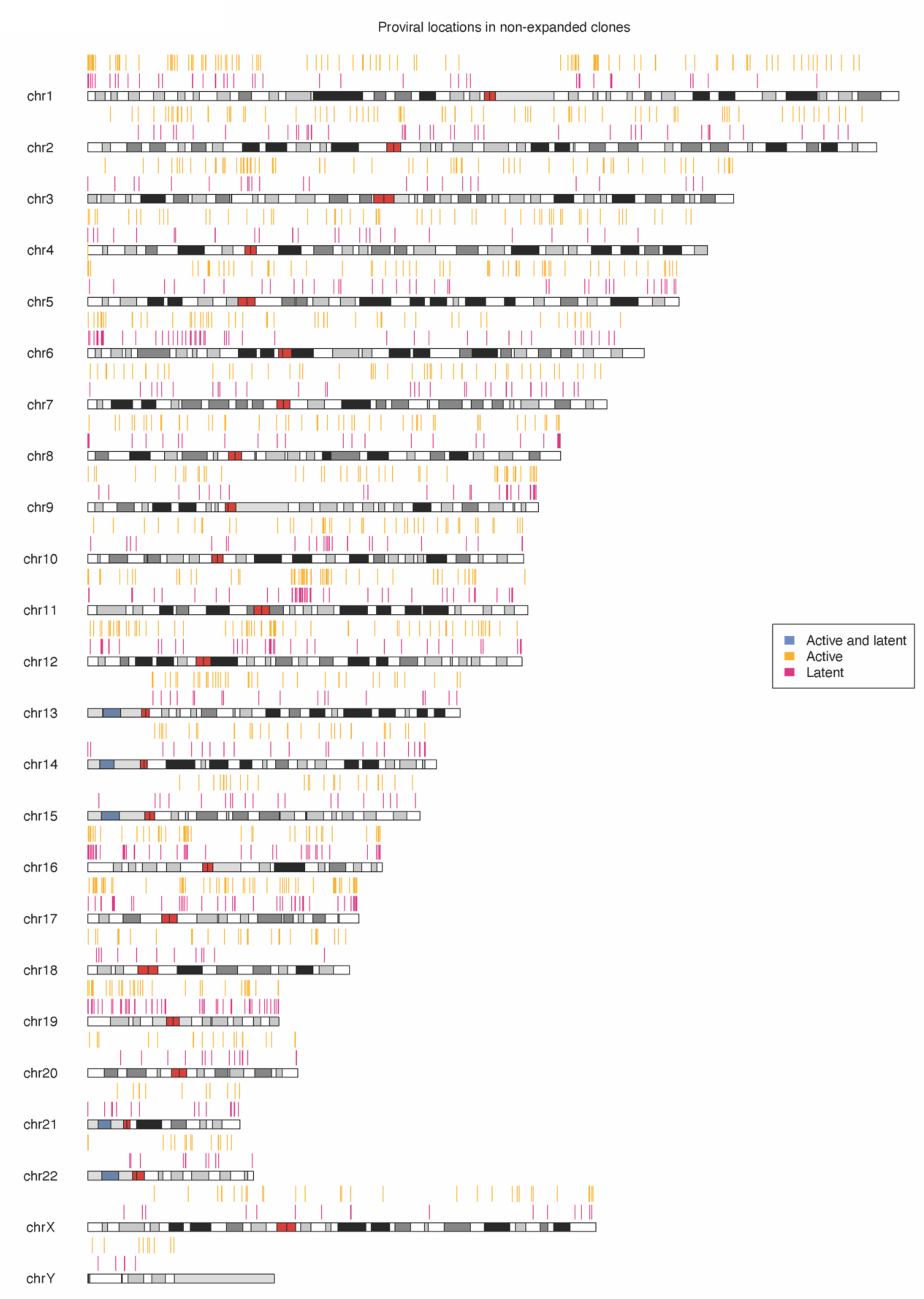
Karyoplot showing the chromosomal distribution of active and latent provirus integration sites identified in non-expanded clones. Each ideogram represents a chromosome, and integration sites are mapped to their genomic coordinates. Proviruses are colored according to transcriptional status, and multiple integrations within non-expanded clones are displayed on the same chromosome. Cytogenetic bands are shown based on Giemsa staining patterns as described for Figure S9.

**Figure S11.**
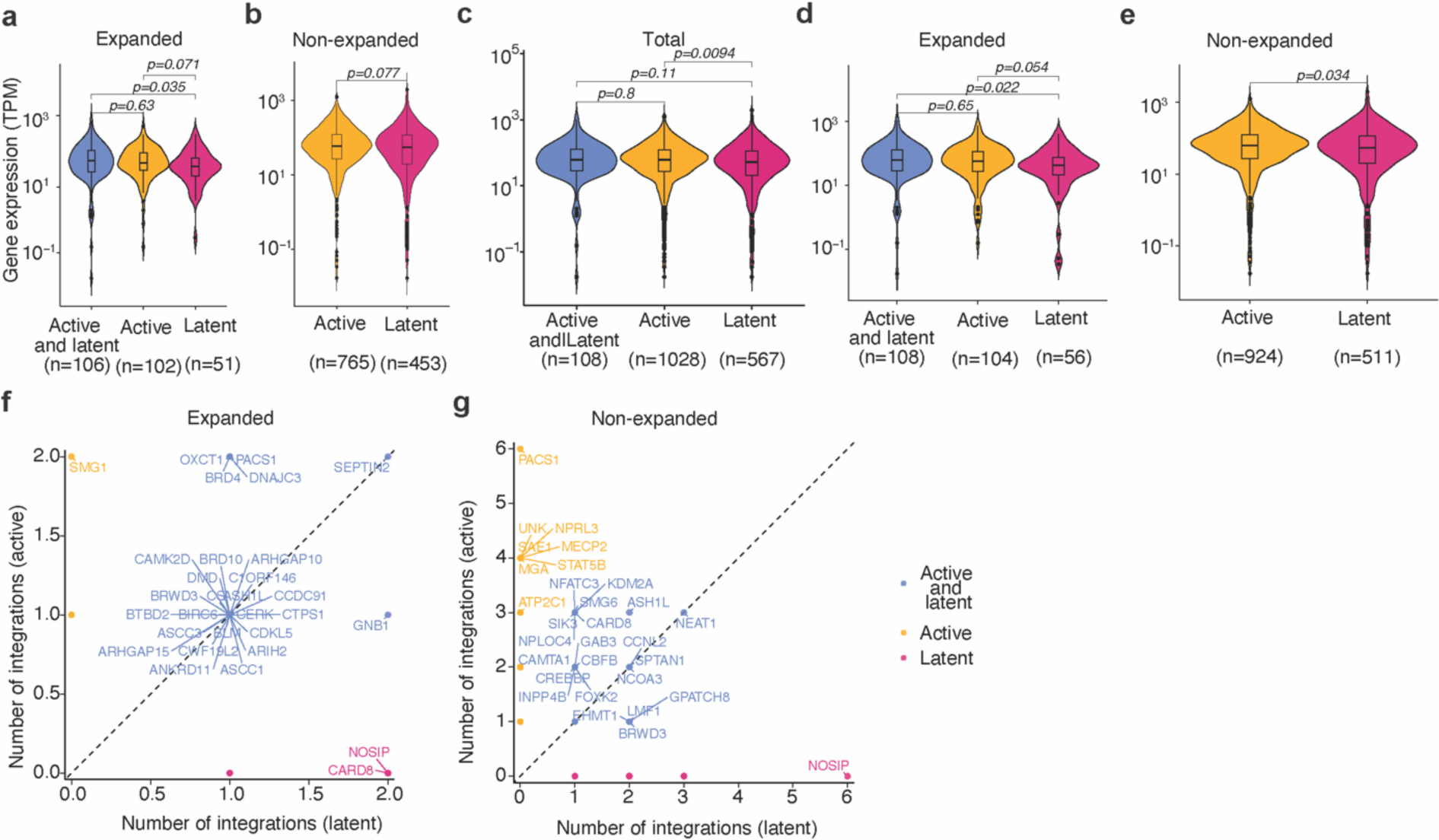
**(a** and **b)** Expression levels of genes with active, latent, or active and latent proviruses for expanded and non-expanded cell clones. **(c-e)** Gene expression associated with individual active, latent, or active and latent proviruses for expanded, non-expanded, and total cell clones. P value denotes the Wilcoxon rank-sum test with Bonferroni correction for multiple comparisons. **(f** and **g)** Number of distinct integrations in genes with active and latent proviruses found in expanded and non-expanded cell clones.

**Figure S12.**
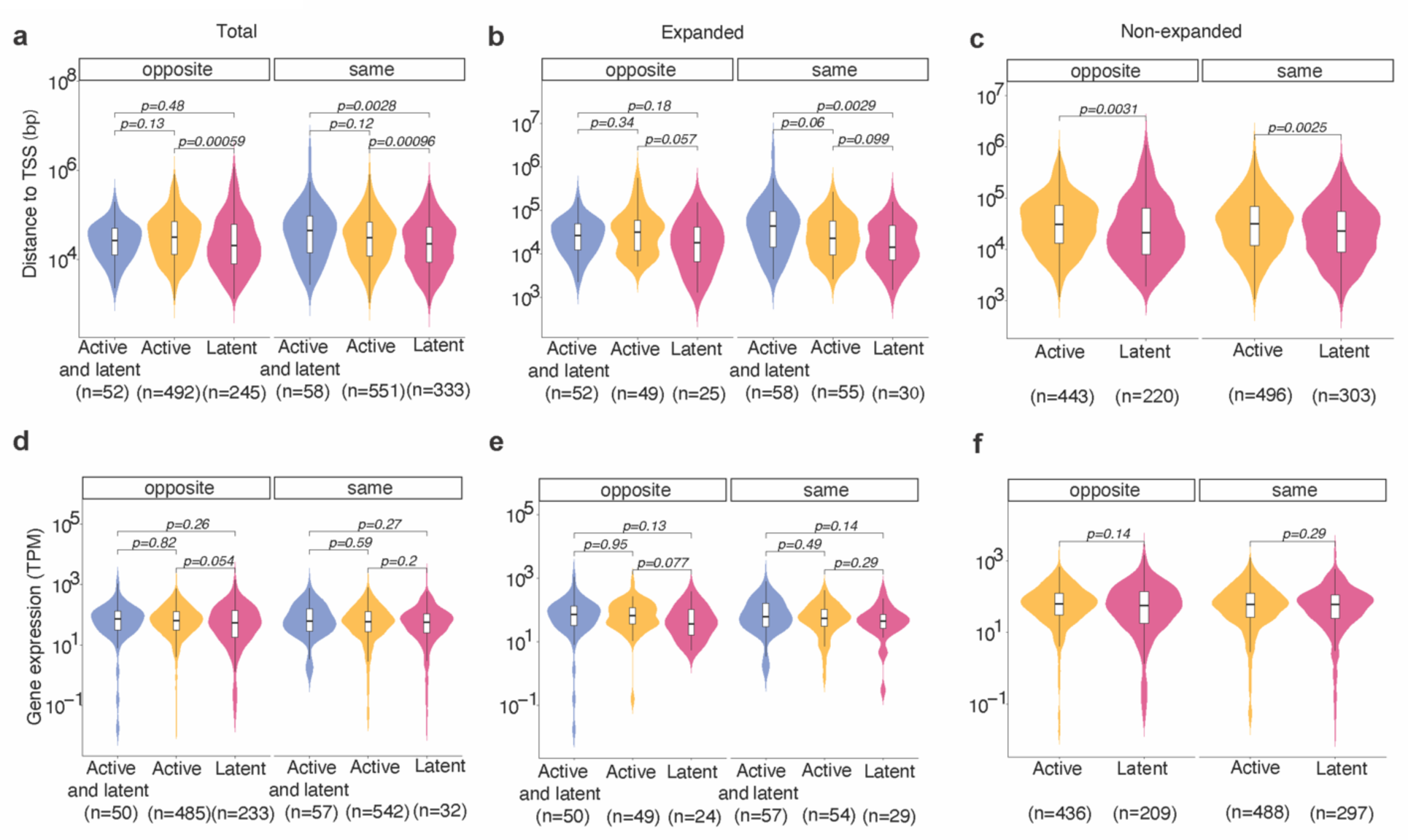
**(a-c)** Distance of integrated genic proviruses from the TSS for proviruses integrated in the same or opposite orientation of the TSS in the expanded, non-expanded, and total cell clones. **(d-f)** Expression of genes with integrated proviruses in the same or opposite orientation with respect to host transcription in expanded, non-expanded, and total cell clones. P value denotes the Wilcoxon rank-sum test with Bonferroni correction for multiple comparisons.

**Figure S13.**
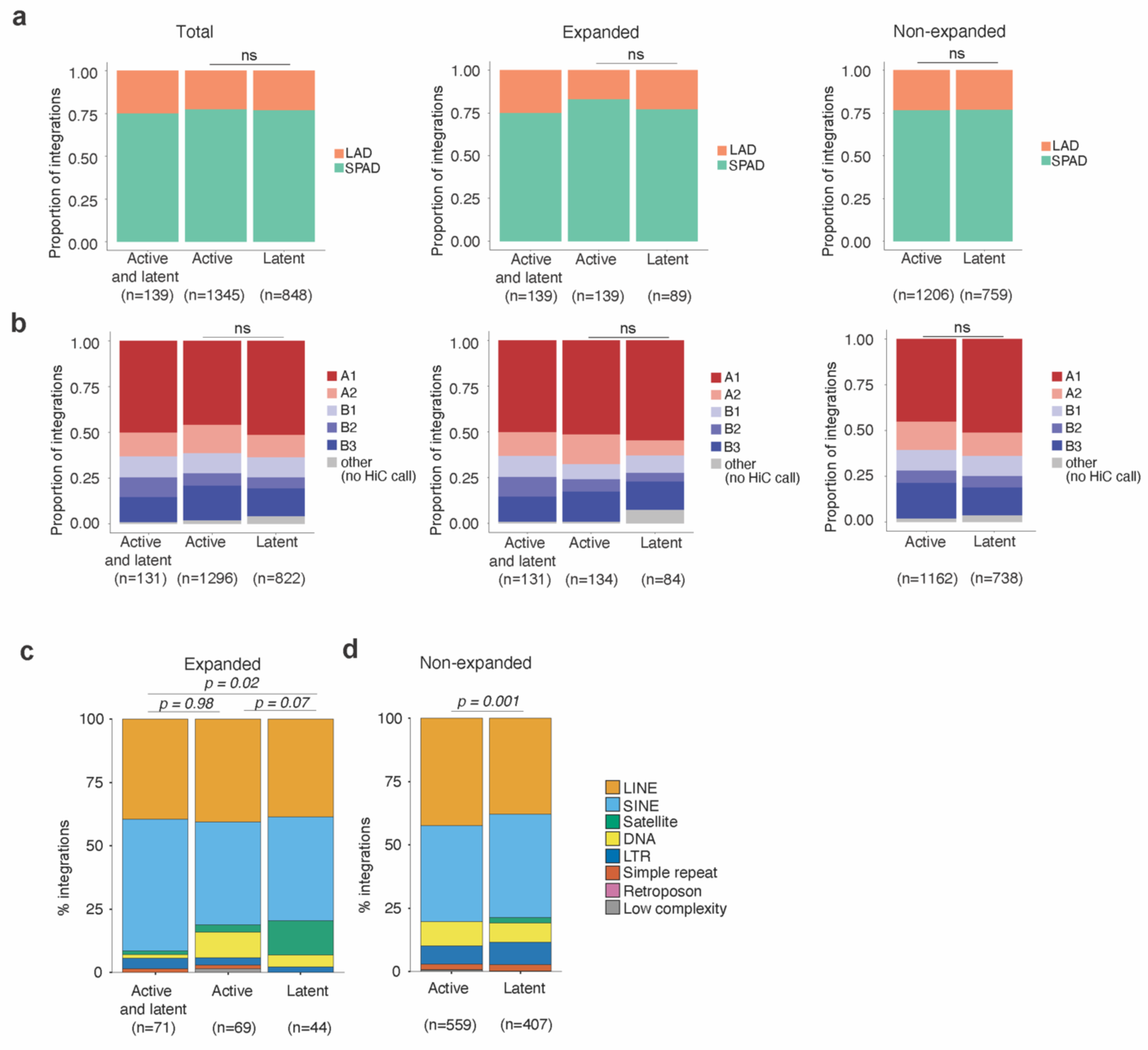
**(a)** Proportion of SPAD and LAD annotation for associated with active, latent, or active and latent proviruses in total, expanded, and non-expanded cell clones. **(b)** Proportion of active, latent, or active and latent proviruses found within chromatin structural compartments A and B and their respective sub-compartments as determined by Hi-C sequencing data for expanded, non-expanded, and total cell clones. P-value denotes the two-proportion z-test. **(c** and **d)** Percent of proviruses in distinct chromosomal repeat regions in the human genome for active, latent, or both active and latent proviruses in expanded and non-expanded cell clones. P-value denotes the two proportions z-test for the satellite repeat elements class.

**Figure S14.**
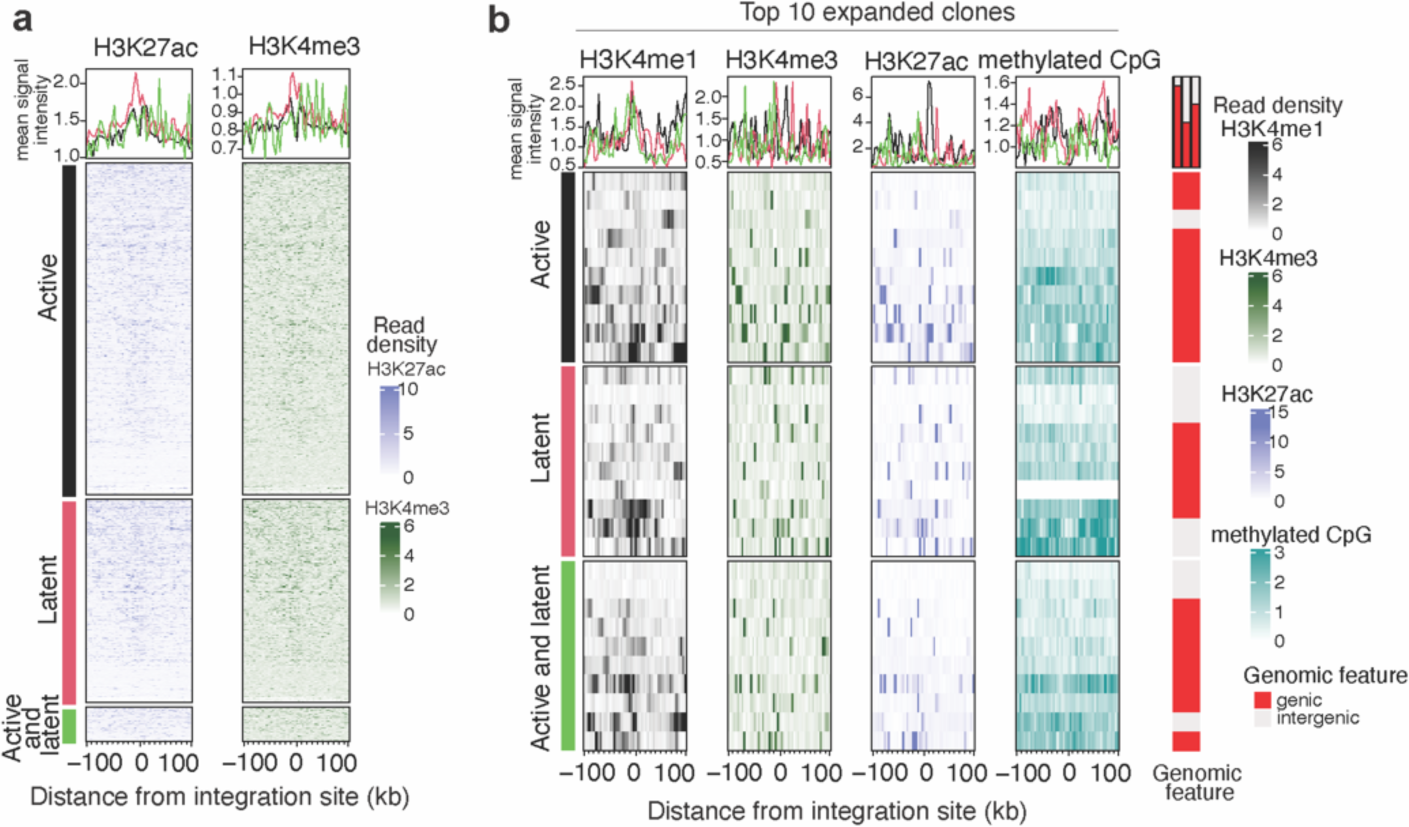
**(a)** Heat map showing levels of histone marks found up to 100kb upstream and downstream of the integration sites of active, latent, or both (active and latent) proviruses. The line plot above the heat map shows the mean signal across all integration sites found within the active (black), latent (pink), and both (green) categories. The heatmaps for each group are arranged from top to bottom according to the highest to lowest mean signal intensity for H3K36me3, as in Fig 5a. **(b)** Heat map showing levels of histone marks found up to 100kb upstream and downstream of the integration sites of active, latent, or both (active and latent) proviruses for the top 10 expanded clones in each category.

**Figure S15.**
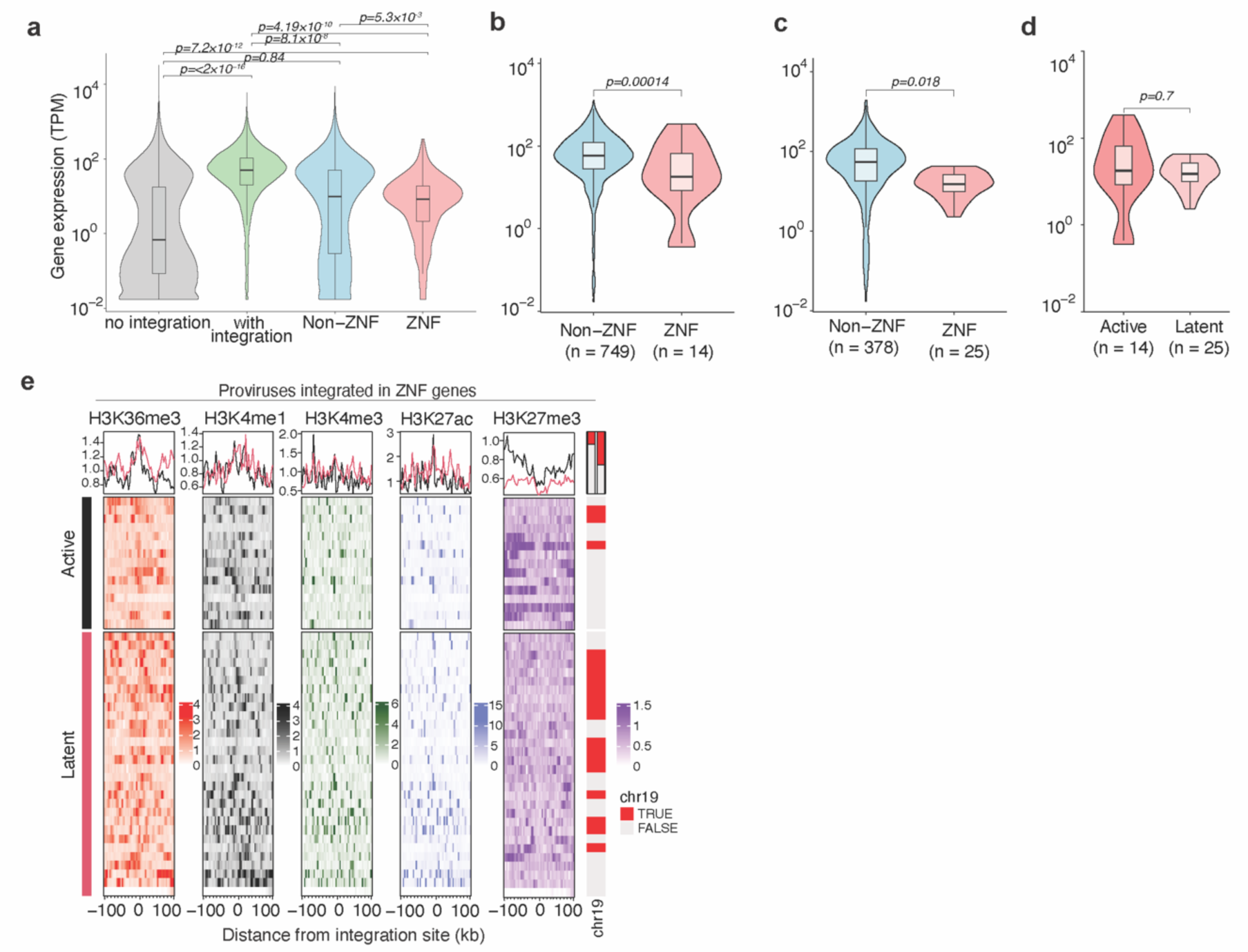
**(a)** Comparison of the level of expression of genes with or without HIV-1 integration, non-ZNF, and ZNF genes. **(b** and **c)** The expression of ZNF genes as compared to non-ZNF genes with either the active (**b**) or latent (**c**) proviruses. **(d)** Comparison of the expression of ZNF genes with active or latent proviruses. The box inside the violin plot shows median, interquartile ranges, and minimum/maximum values. P values were determined by the pairwise Wilcoxon rank-sum test. **(e)** Heat map showing levels of histone marks found up to 100kb upstream and downstream of the integration sites of active, latent, or both (active and latent) proviruses for proviruses integrated in ZNF genes.

